# Convergent evolution of noncoding elements associated with short tarsus length in birds

**DOI:** 10.1101/2024.04.30.591925

**Authors:** Subir B. Shakya, Scott V. Edwards, Timothy B. Sackton

## Abstract

Convergent evolution is the independent evolution of similar traits in unrelated lineages across the Tree of Life. Various factors underlie convergent evolution including convergent rate changes through consistent shifts in substitution rate in the same genes or gene networks. In this study, we use comprehensive phenotypic data to identify seven bird clades with independent shortening in tarsus length and use both comparative genomic and population genetic data to identify convergent evolutionary changes among four target clades with shifts to shorter optimal tarsus length. Using a newly generated, comprehensive set of avian conserved non-exonic elements (CNEEs), we find strong evidence for convergent acceleration in short-tarsi clades among CNEEs, but not protein-coding genes. Accelerated CNEEs in short-tarsi clades are preferentially located near genes with functions in development, with the strongest enrichment associated with skeletal system development. Further analysis of gene networks highlighted this larger role of changes in regulation of broadly homologous developmental genes and pathways as being an integral aspect of limb size variability in birds.

---

Convergent evolution – the independent evolution of similar traits in unrelated lineages – is a common feature across the Tree of Life ^1,2^. Convergent phenotypes have been associated molecular changes at many levels ^2,3^, ranging from shared single nucleotide substitutions (e.g. bird coloration ^4^, snake venom ^5^), to parallel rate shifts in protein-coding genes (e.g., adaptation to marine life ^6^ or echolocation ^7^), to repeated changes in putative regulatory sequences (e.g., *Heliconius* butterfly wing patterning ^8^, stickleback spines ^9^, flightlessness in birds ^10^, and limb loss in squamates ^11^). Understanding the diverse mechanisms facilitating convergent evolution has long been a goal of evolutionary biology.

The recent dramatic increase in the number of high-quality genome assemblies encompassing diverse organisms ^12–14^ has facilitated studies on convergent evolution at every scale. High quality annotations of protein-coding genes from many species ^15,16^, incorporation of ATAC-seq and Chip-seq data ^17,18^, the capacity to generate large-scale whole genome alignments ^19^, coupled with machine learning algorithms to identify regulatory elements ^20,21^ have further aided studies of regulatory changes in convergent evolution. It is, however, important to note that this ‘reverse genomics’ paradigm depends critically on the same genetic loci or gene networks underlying convergent phenotypes; to the extent that when different loci or networks underlie convergent phenotypes, phylogenetic and comparative approaches may be compromised ^22^.

Birds have emerged as an excellent system to study convergent evolution ^10,23^. They have comparatively small, less-complex genomes that are easy to assemble ^24,25^ and a wealth of phenotypic data that can be mapped on dense phylogenies ^14,26,27^. Most studies in avian evolution, however, have been focused on relatively few taxa, and large-scale comprehensive studies of convergent evolution encompassing the full diversity of bird species, and the association of such phenotypes with genomic data have not been systematically performed.

To address this gap, we focus on tarsus length, an easily repeatable, simple linear character in birds that shows a multitude 44of forms, from the long legs of flamingos to the short legs in hummingbirds. Hindlimb size, to which tarsus length contributes, is highly correlated with behavior and natural history and hence, yields a vivid picture of the evolutionary pressures underlying species evolution ^28^. For example, in wading birds, long legs are correlated with depth of foraging to reduce drag while moving across the water ^28,29^. Similarly, aerodynamic and fluid dynamic pressures often lead to shorter legs in aerial and swimming birds ^28^. Such extremes of short and long limb length exist along many diverse groups of birds and, hence, make for a great candidate in a study of convergent evolution.

In this study, we use a comprehensive functional trait database ^27^ covering most bird species to identify clades with convergent shifts in optimal tarsus length, and then identify evolutionary correlates of those shifts to understand the role of convergent molecular evolution in controlling this important ecological trait. Using a new large-scale whole genome alignment from across the bird tree of life, we report a new set of avian conserved elements, annotated with several resources including ATAC-seq peaks, and identify accelerations in these conserved elements correlated with convergent shifts to shorter tarsus length in birds. We also contrasted convergence in acceleration in conserved elements to changes in protein coding sequences to compare roles of protein coding loci and putative regulatory regions correlated with avian short- tarsus phenotype. To complement these comparative genomic tests, we also use population resequencing data from a subset of the short tarsus clades we identify, and test for selection in both protein-coding and non-coding regions of the genome. Together, these analyses provide strong evidence for convergent evolution associated with putative regulatory regions, but not protein-coding genes, in the evolution of tarsus length.

## Results

### Multiple shifts in tarsus length across the avian tree

To identify convergent changes in tarsus length among bird groups, we used AVONET ^27^, a comprehensive functional trait database that reports measurements of tarsus length for all bird species. Using a subset of 5405 measurements of body-size-corrected tarsus length that are represented by genetic data in the all-bird phylogeny ^30^, we fit a multi-state Ornstein-Uhlenbeck model (OU) implemented the program *bayou* ^31^, which detects adaptive shifts in trait optima across a phylogeny. This analysis yielded seven multi-taxon shifts in tarsus length across the bird phylogeny (Fig. 1A, 1B, Supp. Table 1). Two of these shifts were towards longer tarsi, at the node uniting the crab-plover (Dromadidae) and pratincoles (Glareolidae) and at the node uniting the antpittas (Grallaridae). The remaining five were shifts towards shorter tarsi, at the node uniting tropicbirds (Phaethontidae), hoatzin (Opisthocomidae), and sandgrouse (Pteroclidae); at the node uniting penguins (Spheniscidae); at the node uniting kingfishers (Alcedinidae); at the node united swallows (Hirundindae); and at the node uniting bulbuls (Pycnonotidae).

**Figure 1:**
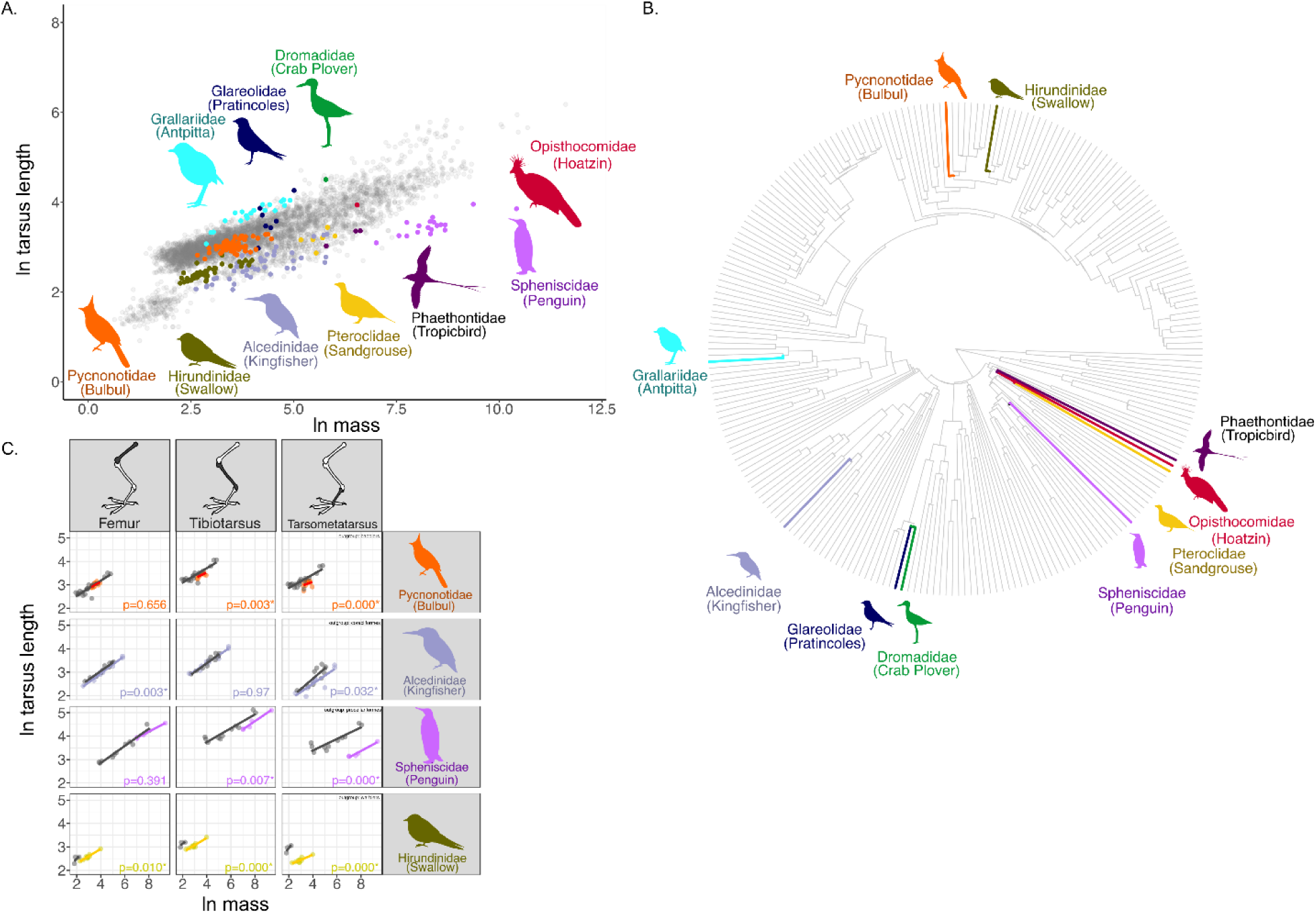
A) Scatterplot of log-transformed values of mass and tarsus length for all birds. Highlighted families were shown to have shifts in tarsus length using an OU model implemented in *bayou*. Individual species where *bayou* also showed shifts are not highlighted in the plot. B) Location of shifts detected using *bayou* placed on phylogenetic tree showing one branch for each of the families of birds. Families showing shifts in tarsus-size are highlighted in the tree. A larger version of the figure with all tip labels is included in the supplemental files. C) Scatterplot of log- transformed values of mass and lengths of femur, tibiotarsus, and tarsometatarsus of short -tarsi shifted families and their respective comparative outgroups. P-values of difference between the measurements of the respective bones in focal group versus outgroups are shown for each comparison. Values less than 0.05 are marked with *.

**Table 1:** Gain of motifs among limb development genes associated with acceleration in conserved elements in the respective target groups.

| ID | Chicken Chromosome | Closest genes | motif | Target group |
| --- | --- | --- | --- | --- |
| ce168790 | 2 | *HOXA9, HOXA10, HOXA11* | Tbx4 | Kingfisher |
| ce246503 | 3 | *MEIS1* | Isl1 | Kingfisher |
| ce442669 | 6 | *SOX6* | Tbx18 | Kingfisher |
| ce1053695 | 24 | *ETS1* | Meis1 | Swallow |
| ce1053695 | 24 | *ETS1* | Fos | Swallow |
| ce1117607 | Z | *SMAD2* | Hoxc12 | Bulbul |

Based on the availability of genomic resources, we chose to focus on four groups well-represented by genomic resources and with shifts towards shorter optimal tarsus lengths: penguins, kingfishers, swallows, and bulbuls. Using skeletal elements from specimens at the Museum of Comparative Zoology, we validated the shifts in tarsus length in all four groups using the tarsometatarsus bone, the skeletal proxy for tarsus measurements in museum skins, which were shorter in all four focal species than in the respective outgroups (p-values, Cohen’s D: bulbul = <0.001, 1.223; swallow = <0.001, 7.142; penguin = <0.001, 2.350; kingfisher=0.032, 0.729; Fig. 1C, Supp. Table 2). We also compared the other two limb bones, femur and tibiotarsus, to test if the entire limb was shorter and found this to be the case in swallows ( p-values, Cohen’s D: femur = 0.010, 4.367; tarsometatarsus = <0.001, 4.835). In penguins and bulbuls, the tibiotarsus was shorter (penguin = 0.007, 1.583; bulbul=0.003, 1.103), but not the femur, while in kingfishers the femur was shortened (0.003, 0.045), but not the tibiotarsus.

### A comprehensive catalog of conserved elements in birds

To generate a comprehensive dataset of annotated conserved elements in birds, we combined two previously published sets of avian conserved elements ^10,32^ with one published vertebrate-wide set of conserved elements (UCSC) and two new avian conserved element sets generated with chicken and zebra finch as reference genomes respectively (Supp. Fig. 1A) using a 79-genome Cactus alignment that also included multiple taxa from all four focal clades (Fig. 2A, Supp. Table 3). All five sets of conserved elements were mapped to the chicken assembly *bGalGal1.mat.broiler.GRCg7b* (Refseq Accession: GCF_016699485.2; subsequently referred to as galGal7b), resulting in a total of 1,117,392 elements with a mean substitution rate of 38% of the neutral rate (mean rho=0.38; Fig. 2B, Supp. Fig. 2). Based on per-base conservation scores computed from a 363-species avian whole genome alignment ^14^, 57.2% of bases were strongly conserved in an average conserved element (Fig. 2C, Supp. Fig. 3). To account for alignment quality issues and missing data, we filtered out elements with less than 30% strongly conserved bases, leaving a final set of 932,467 conserved elements (median length 71 bp, mean length 122 bp, maximum length 4620 bp; Fig. 2B, Supp. Fig. 1A, 4). These elements span the genome but were overrepresented in genes (Fisher’s exact test simulated p < 0.01, odds ratio 1.61): 28.11% of the conserved bases overlap exons, 44.22% spans introns, compared to 26.79% in intergenic regions (Supp. Fig. 5). Upon comparing the conservation status of these elements with those in mammals to assess the phylogenetic specificity, we recovered homologous regions for 35.0% of the elements. Using the Zoonomia 241-species whole genome alignment ^19^, approximately 72% of these homologous elements were also highly conserved in mammals (Supp. Fig. 6).

**Figure 2:**
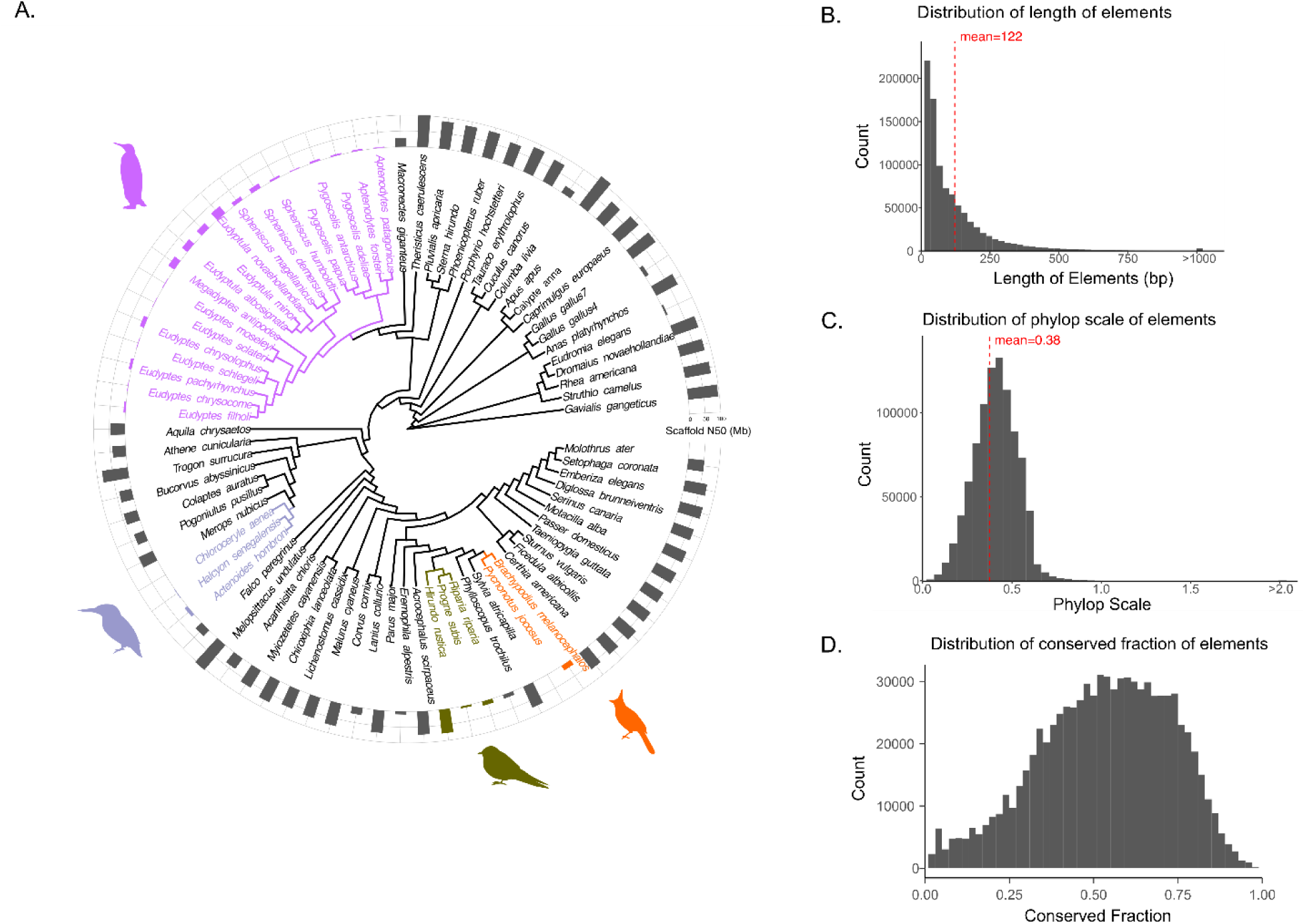
A) Phylogenetic tree showing samples used for Cactus alignment; focal short -tarsi shifted species are represented by respective colors. Bargraph shows N50 scores of genomes highlighting the quality of assemblies used for the Cactus alignment. B) Distribution of lengths of conserved elements. Elements greater than 1000 bp have been collated into one stack. Mean value of elements shown by red line. C) Phylop scale distribution of conserved elements showing the range of the parameter rho, representing the mean substitution rate in conserved elements relative to the neutral rate. Weighted mean value of phylop scale shown by red line. Elements with phylop scale greater than two have been collated into one stack. D) Distribution of fraction of nucleotides that is conserved in each element based on conservation based on per-base conservation scores computed from a 363 species avian whole genome alignment ^14^.

To better understand the putative regulatory function of these conserved elements across birds, we used previously published open-chromatin regions identified by ATAC-seq, which measures accessibility of DNA to transcription factors and is likely correlated with active transcription and regulatory sequence ^18^. We re-called ATAC-seq peaks from multiple replicates of 36 embryonic chicken tissues from eleven published studies (Supp. Table 4) to generate a consistent and comparable set of ATAC-seq peaks predicted on the galGal7b reference genome. ATAC-seq peaks were called using Genrich (https://github.com/jsh58/Genrich), resulting in 860,472 open chromatin regions (mean length 226 bp; Supp. Fig. 7A) representing approximately 18.8% of the genome. Of these, 476,451 (55.37%) were unique to only one of the 36 tissues analyzed, and 5332 were present in all 36 tissues (Supp. Fig. 7B); similar tissue types and closer embryonic developmental stages were more correlated with one another and share more ATAC-seq peaks than dissimilar tissues or developmental stages (Supp. Fig. 8, 9).

Conserved elements were strongly enriched among ATAC-seq peaks (Fisher’s exact test simulated p < 0.01, odds ratio 1.69): 363,838 ATAC-seq peaks (40.43%) overlapped with conserved elements (Supp. Fig. 5).

### Convergent evolution of putative regulatory elements accelerated in short-tarsi clades

To identify patterns of acceleration and convergence in our set of avian conserved elements in short-tarsi clades, we used PhyloAcc ^33^, a Bayesian phylogenetic framework that allows inference of branch-specific shifts in conservation state (Supp. Fig. 1B). We ran three models with PhyloAcc—a null model that does not allow an element to experience acceleration on any lineage; a target model that allows acceleration only in focal lineages (in this case, the species with short tarsi); and a full model that allows acceleration in any lineage—for 726,331 conserved non-exonic elements (CNEES) using respective neutral models generated for macrochromosomes, microchromosomes, and Z chromosome. Using Bayes factors to compare models, we identified 14,422 CNEEs where the target model is supported over the null model by a Bayes factor of at least 10 (Supp. Fig. 10), implying strong evidence for acceleration in at least one short tarsi species. These accelerated elements had a mean conserved rate 0.26 times slower and a mean accelerated rate 1.91 faster than the neutral rate (Supp. Fig. 11). To exclude idiosyncratic acceleration in a single species, we focused on the 7948 CNEEs that are accelerated in all species within the relevant focal clade(s) (or more than 3 penguins), which we subsequently refer to as the short-tarsi broad dataset (Supp. Fig. 1C). We also identified a subset of 403 accelerated elements where the target model is supported over the full model by a Bayes factor of at least 5, representing elements with evidence for acceleration exclusively in short-tarsi species (the short-tarsi specific dataset subsequently; Supp. Fig. 1C).

Short-tarsi accelerated elements were convergently shared between the four focal groups. Using hypergeometric tests, we found evidence for more shared acceleration in the short-tarsi broad dataset between all pairwise groups of swallow, kingfishers, and penguins than expected by randomness (p values-swallow vs kingfisher: 1.46 x 10^-9^, swallow vs penguin: 7.55 x 10^-2^, penguin vs kingfisher: 1.68 x 10^-167^), although not between bulbuls and other groups. In the short-tarsi specific dataset, we found evidence for shared acceleration between bulbul and penguin (6.88 x 10^-3^), penguin and kingfisher (3.71 x 10^-2^), and penguin and swallow (6.67 x 10^-2^). Using comparable outgroups for each of the four clades, we found evidence for higher number of shared accelerated elements in the short-tarsi broad dataset between all four clades (Fisher’s p-value < 0.01), between three-way comparisons of bulbul, swallow, and penguin (< 0.01) and between bulbul, kingfisher, and penguin (< 0.01), and two-way comparisons of bulbul and kingfisher (0.03), bulbul and penguin (< 0.01), and kingfisher and penguin (< 0.01) (Fig. 3A).

**Figure 3:**
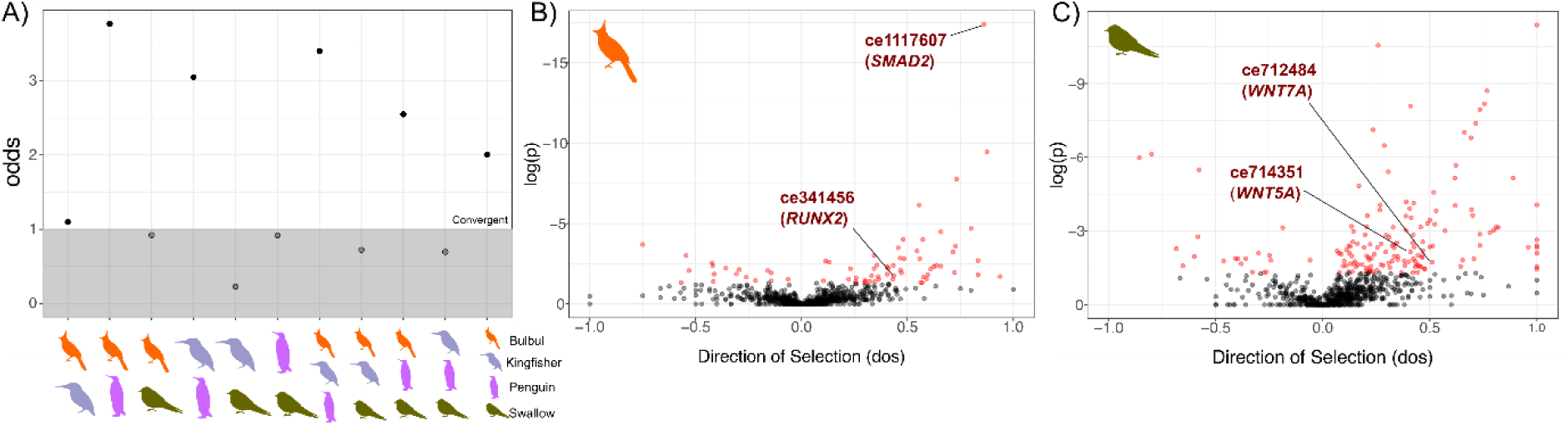
A) Test of convergence for shared accelerated elements between four focal groups, bulbuls, kingfishers, penguins, and swallows using comparative outgroups. Likelihood odds from Fisher’s exact test for each comparison are shown with points above grey box being significantly enriched in the respective target groups than the respective outgroups. B) Direction of selection from a modified McDonald & Kreitman test for conserved elements, compared with the synonymous sites in the closest gene, among the short-tarsi broad dataset that are accelerated in bulbuls and B) swallows. For bulbuls, population-level whole genome resequencing data for *Brachypodius melanocephalos* was used while for swallows, data for *Hirundo rustica* was used. Significant values for elements are colored red. A few elements with genes close by that are relevant for limb development are highlighted in each plot.

### Evidence of positive selection on short tarsi accelerated elements

For two focal groups with publicly available population level whole-genome resequencing data, bulbuls ^34^ and swallows ^35^, we also evaluated evidence for selection. Using a modified McDonald-Kreitman test ^36,37^ comparing fixed and polymorphic sites in accelerated elements to those in synonymous sites in the closest protein coding genes, we found 54 elements showing positive selection and 17 elements showing relaxed constraint in bulbuls (Fig. 3B) and 138 elements showing positive selection and 20 elements showing relaxed constraint in swallows (Fig. 3C). While we did not recover any significantly enriched GO terms, several elements near limb development genes that have been implicated in short limb phenotypes were found to be positively selected. In bulbuls, two positively selected elements were present near *SMAD2* and *RUNX2. SMAD2* facilitates downstream Tgf expression, misregulation of which has been shown to lead to shorter limbs in rats ^38^. Dosage changes of *RUNX2* has also been shown to reduce limb size through its effect on cartilage development ^39,40^. Similarly, two Wnt genes important for limb development, *WNT5A* and *WNT7A,* had conserved elements nearby that were positively selected in swallows. Loss of function of *WNT5A* is known to result in short limbs as it leads to an inability of extension of the AP-axis ^41^. *WNT7A* performs multiple functions in the developing limbs and misregulation can lead to various defects of limb development ^42^.

Since each conserved element is relatively short, we also evaluated evidence of selection in a combined set of all conserved elements within 10 kb of each gene (referred to as CE cluster), including elements within intronic regions of the gene. Using an alpha selection cutoff of 0.5, we found 104 positively selected CE clusters in bulbuls (Supp. Fig. 12A) and 263 in swallows (Supp. Fig. 12B). No GO term was found to be enriched in genes near CE clusters in either bulbuls or swallows. However, among the 19 CE clusters that were positively selected in both bulbuls and swallows, we identified one near the limb development gene *SHH*. *SHH* is an important limb development gene with many functions in limb formation and limb patterning ^43^, and could have a role in limb shortening.

### Short tarsi accelerated elements are associated with open chromatin and limb development genes

Short tarsi accelerated elements were more likely to be within open chromatin regions than a randomly sampled conserved element (Fisher’s T-test p-value: < 0.01, odds ratio: 1.98). Of the 7948 accelerated elements in the short-tarsi broad dataset, 4151 (52.2%) overlapped chicken ATAC-seq peaks of which 1575 were open in chicken hindlimbs at HH18 stage while in the short-tarsi specific dataset, 216 of the 403 elements (53.5%) overlapped ATAC-seq peaks with 102 elements open in hindlimbs at HH18 stage. Since the ATAC-seq peaks were based on chicken experiments, and not the target species, to account for possible variation in chromatin openness across the phylogeny, we used TACIT ^20^, a convolutional neural network-based machine-learning approach, to train a classifier allowing us to predict the probability of elements being open in the other species. Using the ATAC-seq peaks called for the hindlimb tissue at the HH18 stage for chicken and emu ^44^ as the training dataset to predict ATAC-seq peaks in all species, we found, in the short-tarsi broad dataset, 110 elements that were open in the chicken but predicted as closed in the accelerated lineages and 99 elements where the chicken state was closed but was predicted to be open in the accelerated taxa (Supp. Table 5). While functional enrichment test of GO terms of genes near these elements yielded no enriched GO categories, some elements near limb development genes like *ETS1* and cartilage development genes like *SOX5* and *SOX9* had shifts in openness in our short-tarsi target lineages.

Short-tarsi accelerated elements were specifically enriched for genes that were involved in skeletal and limb development. Using PANTHER to categorize GO (Gene Ontology) biological process terms associated with 5008 and 445 genes within 10kb from the short-tarsi broad and specific dataset respectively, we found 30 and 80 GO categories that were statistically overrepresented, while accounting for the nonrandom distribution of CNEEs in the genome (see Methods for details; Supp. Table 6). Based on semantic similarity between terms (as calculated by *simplifyEnrichment* ^45^), the biological processes enriched in the broad dataset include appendage development, nitrogen response and cellular transport (Fig. 4A), while the biological processes enriched in the specific dataset include regulation, appendage development, organization, Wnt signaling, and stem cell proliferation (Fig. 4B). To identify genes and pathways most relevant to the short tarsi phenotype, we looked specifically for functions that are more strongly enriched in the short-tarsi-specific dataset than the short-tarsi-broad dataset.

**Figure 4:**
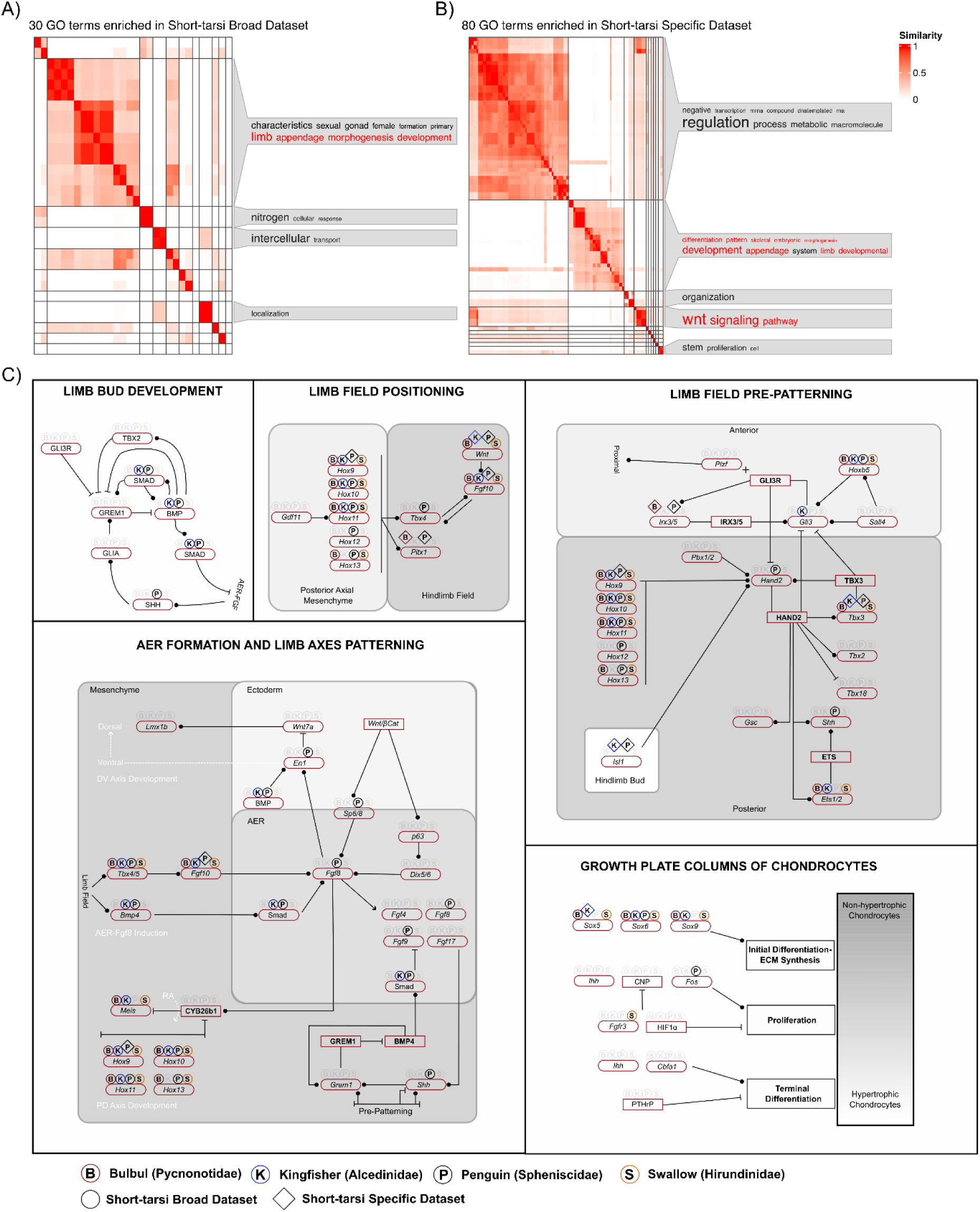
Heatmaps showing clustering of enriched GO categories in the A) short-tarsi broad and B) short-tarsi specific dataset. Word clouds show the most common terminology in the description of clustered GO categories. Categories in red represent terminology relevant to limb development. C) Schematic of genes involved in early hindlimb development—limb bud development, limb field positioning, limb field pre-patterning, and AER formation and limb axes patterning—adapted from Zuniga and Zeller (2020) along with genes involved in growth plate formation in chondrocytes. B, K, P, and S on top of genes refer to whether a gene has a conserved element nearby that is accelerated in bulbuls, kingfishers, penguins, and swallows respectively. Rounded boxes represent genes and square boxes represent proteins/transcription factors. Direction of interaction is denoted by circular ends to represent upregulation and straight perpendicular line to represent inhibition. Abbreviation in circles represent accelerated elements that are part of the short-tarsi broad dataset while those in diamonds short-tarsi specific dataset. Multiple elements may be near any one gene.

Eleven GO categories showed dramatic increases in enrichment in the short-tarsi specific dataset that included three—embryonic skeletal system development, chondrocyte differentiation, and forelimb morphogenesis—that are related to limb development (Supp. Fig. 13).

### Gene-level regulatory convergence in genes with known functions affecting limb

While there were significantly enriched GO terms for genes near accelerated CNEEs that related to skeletal system development, GO categories can be quite generalized and often incomplete and biased ^46^, so we additionally looked for accelerated CNEEs in known pathways linked to limb development (Fig. 4C, Supp. Fig. 14). We found multiple clusters of genes relevant to skeletal and limb development near accelerated elements (Fig. 4C), many of which show evidence for convergent acceleration at the gene level (defined as accelerated CNEEs near that gene in multiple focal taxa, regardless of whether the specific accelerated CNEEs are the same or different among focal taxa).

Two genes are particularly notable: *FGF10* and *PITX1,* which are both active in early embryonic limb development through limb field positioning (Fig. 4C) ^43,47^. Eleven elements near *FGF10,* an integral protein required in early limb development, the misregulation of which can result in severe truncation of limbs ^48,49^, were accelerated across all four focal groups. *PITX1,* with 2 nearby elements accelerated across penguins and bulbuls, is a hindlimb specific homeodomain factor, which can cause long bones in the hindlimb to be shortened if altered ^50^.

Another gene, *MEIS1,* overexpression of which affects proximodistal limb axis development ^51^, was near 9 elements accelerated across bulbuls, swallows, and kingfishers.

Hox genes, *SOX9* (which can lead to short limbs through its activity at multiple stages of chondrocyte differentiation ^52,53^) and two Wnt pathway genes (*WNT5B* and *WNT11,* both facilitating chondrocyte differentiation ^42^) also show evidence for gene-level convergent acceleration. Two elements near *SOX9* and are accelerated in bulbuls, kingfishers, and penguins while *WNT5B* is accelerated in kingfishers and penguins and *WNT7B* accelerated in bulbuls and kingfishers. Among Hox genes, convergently accelerated elements were present close to *HOXA9* as well as a *HOXB* and *HOXD* cluster. Hox genes, especially from the 9-13 group, are expressed along proximal and distal limb elements and facilitate proximodistal patterning different regions of the hindlimb ^54–57^. Hox genes also act in a dosage-dependent manner and the regulation of Hox genes especially groups 10 and 11 can have highly significant effects on limb size and development ^58^. Hox genes, however, tend to be clustered together in the genome ^59^, such that multiple Hox genes are near multiple conserved elements, which prevents us from identifying specific Hox genes that are associated with a particular element.

### Species-specific motif changes in accelerated elements

A few accelerated elements showed species specific gains in motif binding sites in genes relevant for limb development. Using MAST to annotate motifs for genes in known limb development pathways ^43^ in accelerated elements of the focal groups, we found five potential gains of new motifs in specific focal lineages (Table 1). Two novel motifs for Meis1 and Fos were detected in an accelerated element near *ETS1* in swallows. While these genes are integral for limb development, we did not find any previous work suggesting direct interaction between these genes. This element overlapped with a chicken ATACseq peak but was predicted to be closed in swallows, suggesting changes in chromatin accessibility at development. We also recovered a potential Tbx4 motif in an accelerated element among Hox cluster a9-a11 and a Tbx18 motif in an accelerated element near *SOX9* in kingfishers. TBX4 has been shown to interact with HOXC11 and HOXD11 and binding to T-box-Hox composite DNA motif that then show synergistic activity in downstream processes ^47,60^. *TBX18* is expressed in early limb development and facilitates somite maturation especially in the stylopod portion in the limbs ^61^. *TBX18* and *SOX9* interactions have been observed in the development of the urinary system in mice ^62^ but their specific interaction in limb development is unknown. Similarly, we found a Hoxc12 motif in an accelerated element near *SMAD2* in bulbuls. Smad is known to downregulate Hox expression, which then affect the downstream expression of Bmp genes ^63^. This element, ce1117607, also showed a significant value of positive selection (dos = 0.86) in the modified MK test for bulbuls (Fig. 3B) suggesting an active functional change and subsequent selection in bulbuls. While the specific Hox interactions that we observed have not been reported, due the redundancy of paralogous Hox genes, these alternate Hox genes could facilitate similar morphological variation ^54,57^.

### Positively selected genes do not show convergent selection in short-tarsi focal groups

Using multiple approaches, we failed to find evidence of convergent evolution of protein coding sequences in short-tarsi focal clades, in contrast to extensive evidence for convergence in putative regulatory elements (CNEEs). We recovered 9854 one-to-one orthologous loci using TOGA with subsequent filtering for quality and number of taxa. Using aBS-REL ^64^, a branch-site based test of selection that accounted for synonymous rate variation ^65^, we identify 1534 genes with selection in least one short-tarsi focal clade. While pairwise hypergeometric tests for each of the four lineages showed evidence for more shared genes showing selection than random among genes in the four focal groups except for between swallow and bulbul, and between bulbul and kingfisher (Supp. Table 6), Fisher’s exact test comparing genes under selection in the focal group and in representative outgroups showed no evidence of convergence for any pairwise, three-way, or four-way comparisons (Supp. Table 6). GO enrichment analysis of these 1534 genes also did not recover any significantly enriched GO category, suggesting little evidence for convergence at the pathway level.

Assessing selection using the population resequencing data for bulbul and swallow by applying the MacDonald-Kreitman test ^36^, which compares synonymous and non-synonymous polymorphism to divergence in order to identify genes with an excess of non-synonymous fixations, modified to account for the likelihood that rare non-synonymous polymorphisms may represent weakly deleterious mutations ^66^, we fail to detect evidence for convergent positive selection. We found 1069 loci positively selected in bulbul and 2584 in swallow (Supp. Fig. 15). Of these 322 selected genes were shared between bulbul and swallow. The number of shared genes was not significantly higher than expected by chance (p=1.0). In bulbuls, 21 GO terms were enriched and in swallows, 75 GO terms were enriched but none of the terms were relevant to limb or skeletal development. These results are consistent with previous studies showing relatively little evidence for convergent signals in protein-coding genes associated with skeletal morphology ^10^.

## Discussion

Shorter tarsi are common in highly aerial birds (birds spending a considerable proportion of their time on the wing), hanging, or climbing birds, and swimming species ^28^. The six groups we found with shifts to short tarsi follow the predictions from ^28^ with swallows, an aerial insectivore; pratincoles and tropicbirds, aerial feeding sea birds; and kingfishers and penguins, two diving and plunging species. Sandgrouse, an arid adapted group of birds, have short tarsi potentially as an adaptation to living in xeric environments ^67^. The short tarsi in hoatzin, a species with unusual locomotory strategies in juveniles ^68^, may be driven developmental constraints of this behavior. Bulbuls, an insectivorous and frugivorous group of passerine birds found throughout Asia and Africa, do not neatly fall into any category of adaptive explanation for shorter limbs. However, taxon names such as *Microtarsus, Micropus, Hemitarsus, Brachypodius,* and *Brachypus* have been applied to many bulbul species ^69^ indicating their inclusion among short-tarsi birds is expected. One species-rich aerial insectivore group, the Strisores (swifts, hummingbirds, nightjars, and allies) is noticeably absent in our results for short tarsi. This result is probably a caveat of the approach we used, which involved splitting the dataset into multiple superclades and running independent runs of *bayou,* a practice that prevents us from detecting shifts occurring towards the base of the tree. Even without the Strisores, the other groups in our analyses include a well-supported list of birds known to have short tarsi.

Our results support previous studies of convergence in limb morphology suggesting convergent evolution in regulatory elements putatively associated with genes in the same developmental pathways facilitate changes in limb-related phenotypes. In palaeognaths, convergent evolution of forelimb shortening was correlated with changes in rates of nucleotide substitutions in regulatory elements of broadly homologous developmental genes and pathways, albeit often involving different regulatory elements ^10^. Similarly, limb variation in *Anolis* lizards, a clade in which multiple shifts in limb length have occurred, involves different elements near concordant early development pathways that influence limb length ^70–72^. In our study, we found multiple genes along the same limb development pathways with accelerated elements in different genes in different focal groups; some elements additionally show evidence for positive selection or new DNA-binding motifs of genes involved in limb development in specific focal taxa, supporting the importance of pathway level convergence.

In many morphologically convergent phenotypes, especially skeletal traits, regulatory elements undergoing convergent acceleration show more functional coherence than do positively selected protein coding loci for convergent phenotypes ^73^. While we find limb development pathways are significantly enriched among accelerated elements, tests of positive selection in protein-coding genes do not yield any overrepresented function relevant to limb development. Although some overlap between positively selected genes and genes near accelerated elements are expected, and changes in genes themselves probably do play some role in the short limb phenotype, our results show a much larger signal for the role of regulatory regions than protein-coding genes.

Our results suggesting a strong association of convergence in shifts to short tarsi in birds to accelerated rates of substitution in conserved elements in genes near limb development pathways can be utilized for future developmental studies to validate the role of these elements in limb development. This study also provides an annotated, comprehensive list of conserved elements that can be utilized in many future studies on convergent evolution in birds. The nearly one million elements provided in this study can be lifted over to any other genome and hence will be useful for the scientific community in general to use as a basis of their studies. The inclusion of chicken embryonic ATAC-seq peaks can be used to enhance further studies to understand the developmental role of the noncoding regulatory elements in-vitro or in-vivo. The general approaches we applied and resources we provided in this project can be applied to a diverse array of phenotypes and help better understand the evolutionary toolkit underlying phenotypic diversity across the Tree of Life.

## Methods

Briefly, our main goals were to identify convergence in limb-size change in birds and use a newly generated comprehensive annotated list of conserved elements to find acceleration in conserved non-exonic elements (CNEEs) that are correlated to changes in limb size. We identified such shifts among the entire avian lineage using a reversible jump multi-optima OU model and then used the list of conserved elements to find evidence of acceleration in substitution rates using the PhyloAcc framework ^33^. We further investigated the genes around accelerated CNEEs to identify potential candidate genes and pathways that are associated with shifts in tarsus length in birds. All code used in this project is available as RMarkdown files in the supplement.

### Assessing convergence in shifts to short-tarsi in birds

#### Locating shifts in tarsus length

To locate shifts in tarsus lengths in birds, we used a reversible jump multi-optima OU model implemented in a Bayesian framework using the program *bayou* ^31^. The tarsus length data was obtained from AVONET ^27^ along with the phylogenetic tree from ^30^ that was used and modified by ^74^ with tips mapped between phylogenetic tree and morphological traits. The tarsus length values for 5405 species were log-transformed and used as input of *bayou.* We also included log-transformed values of body weight for each species as a correlate in the *bayou* model. Because of the high computing-time requirements to run these models in very large trees, we used an approach similar to ^74^ to divide the dataset into smaller clades. The full dataset was divided into 18 clades (with 19 to 781 species; Supp. Table 1C). As mentioned in ^74^, one limitation of this approach is that shifts at the base of these clades cannot be explicitly tested. However, for the purposes of this project, we decided to focus only on shifts within these clades.

Our priors for *bayou* follow similarly to those used by ^74^ . For each clade, we set a Poisson prior distribution on the number of allometric shifts (*dk*). We used a minimum *λ* parameter of 0.05 times the number of species (2%) in that clade rounded up to the nearest integer or a value of 1 if this number was less than 1. For the maximum *λ* parameter, we used a value of 0.5 times the number of species (50%). We used a uniform prior on the probability of location of shift over all branches on the phylogeny (*dloc*). We set half-Cauchy prior distributions on *α* and *σ*^2^ (*dalpha* and *dsig2*) with a scale parameter of 0.1. For slopes (*dtheta*), we used a normal distribution with a mean of 0.3 and a standard distribution of 0.5 based on the assumption of cubic isometry of body weight and tarsus length. For intercepts (*dbeta_lnMass*), we also used a normal distribution with a mean value of 2 and standard deviation of 0.5. For each clade, we ran short 50,000 generation runs to adjust tuning parameters. We updated prior values of *α*, *β*, *σ^2^* and *θ* to get the acceptance ratio between 0.2 and 0.4.

For each clade, we ran two parallel runs for a minimum of 20,000,000 generations, sampling every 100 generations with a burn-in of 50%. Convergence of runs was checked by calculating Gelman and Rubin’s R using the *gelman.R* function in *bayou* for all estimated parameters. For runs where the effective sample size (ESS) was less than 200 or R was greater than 1.05, we continued the MCMC run for 10,000,000 generations. We continued this process until the ESS was greater than 200 and R was less than 1.05 or we reached 50,000,000 generations, because of the high computing resources required. For a few runs, particularly those with more than 700 species, we did not reach convergence in some parameters, especially if the different runs had stabilized on different number of shifts but we kept the shifts that were present in both runs (Supp. Table 1A, 1B). To identify shifts, we compared the posterior probability between the two runs and inferred shifts in branches that had similar posterior probability scores greater than 0.1. In cases where we did not see convergence among runs, we only chose shifts that were common between the two runs.

We focused on groups that had shifts to shorter tarsus lengths. In order to determine whether only the tarsus was shorter or if all three limbs of the hind bone were shortened, since different mechanisms might underlie the different phenotypes, we measured the length of the femur, tibiotarsus, and tarsometatarsus bones of representative species of the four target lineages and their sister group available at the Museum of Comparative Zoology (Supp. Table 2). For the tibiotarsus, the length to the top of the cnemial crest was not included in the measurement of penguins and Procellariformes as the Procellariformes had highly extended cnemial crests. To avoid measurement bias, SBS measured all the bones using the same calipers, except for the penguins and Procellariformes, which were measured using a larger set of calipers to account for the much longer bones. In total, we measured the three bones for 7 penguins and 11 tube-nosed birds (Procellariformes), 11 kingfishers and 12 non-kingfisher Coraciiformes, 10 swallows and 3 warblers, and 12 bulbuls and 20 babblers.

### Assembling a comprehensive list of conserved elements

#### Genome alignments

79 genomes (Supp. Table 3) were selected for alignment using Progressive Cactus ^19^. The selection of genomes was based on several criteria. We included multiple species of the four target lineages and complemented that with genomes of taxa interspersed across the avian tree of life for background. We also selected only high-quality genomes with most genomes having a scaffold N50 greater than 10 Mb. Two genomes from *Gallus gallus, bGalGal1.mat.broiler.GRCg7b* (GCF_016699485.2; henceforth referred to as galGal7b) and galGal4 (GCA_000002315.1), were used for backward compatibility with previous alignments that used galGal4 as the reference genome. A few genomes of target group taxa were included even if they did not meet the criteria for high-quality. The genome of a crocodylid, *Gavialis gangeticus* (GCA_001723915.1), was included as the outgroup to birds. Most genomes were published genomes from NCBI. For three species, *Struthio camelus*, *Dromaius novaehollandiae*, and *Rhea americana*, we included newly scaffolded genomes from DNAZoo (https://www.dnazoo.org/assemblies). The genome for *Macronectes giganteus* was downloaded from http://genome.kusglab.org/ ^75^.

All genomes were already soft-masked for repetitive elements except for the *Macronectes giganteus* genome. For that species, we used RepeatModeler v2.0.3 ^76^ to generate repeat library.

Then we used RepeatMasker v4.0.6 to first mask repeats using the native Aves repeat library and then the repeat library generated from RepeatModeler. For *Brachypodius melanocephalos*, we supplemented the published genome (GCA01339861.1) with scaffolds from the Z chromosome that were missing in the original assembly ^34^ using the approach mentioned in ^77^, which was similarly lacking Z chromosome scaffolds as a potential consequence of assembly process by Dovetail Meraculous contig assembly pipeline. For all genomes we removed scaffolds that were smaller than 1000 bp to prevent Cactus runs from failing due to their small size.

For the guide tree needed for progressive cactus, we used the tree from ^14^ where we kept only the tips for the species selected for alignment. We added tips that were missing in the tree file but are present among the selected taxa for genome alignment. The *Gavialis gangeticus* tip was appended to the outgroup. Tips for *Eudyptula novaehollandiae, E. robustus* and *E. albosignata* were appended to the tip with *E. minor*. The tip for *Eudyptes schlegeli* was appended to *E. chrysolophus.* For *Lichenostomus cassidix, Corvus cornix, Brachypodius melanocephalos,* and *Emberiza elegans,* the tips for *Lichenostomus melanops, Corvus corone, Brachypodius atriceps,* and *Emberiza buchanani* respectively were substituted. The node for *Gallus gallus* was duplicated to accommodate the two chicken genome assembly versions.

We used the gpu based implementation of progressive Cactus v2.1.1 to align the genomes.

We used the snakemake based implementation of cactus from https://github.com/gwct/GenomeAnnotation/tree/master/genome_alignment to set up the cactus alignment. The Harvard Cannon Supercomputing Resources were used for the run.

#### Generating list of conserved elements

We used a combined approach of using available lists of conserved elements and generating a new list of conserved elements to come up with a comprehensive list of conserved elements. All elements were mapped to the *Gallus gallus* genome galGal7b. Previously available list of conserved elements come from an evaluation of conserved elements in palaeognathous birds ^10^ (henceforth referred to as ratite dataset), conserved elements from vocal learning birds ^32^ (henceforth referred to as the vocal dataset), and conserved elements from a 100-way alignment of vertebrate genomes (http://genome.ucsc.edu/index.html; henceforth referred to as the UCSC dataset). The conserved elements for ratite dataset were lifted over from the galGal4 genomic coordinates (GCF_000002315.3) to the galGal7b genomic coordinates using the NCBI Genome Remapping Service (https://www.ncbi.nlm.nih.gov/genome/tools/remap). The lifted over file was cleaned using bedtools merge to join fragmented regions and then filtered for elements less than 20 bp and greater than 5000 bp. For the vocal dataset, the three individual list of conserved elements from hummingbirds, parrots, and oscines were lifted over from the galGal5 genomic coordinates (GCF_000002315.4) to the galGal7b genomic coordinates. The three datasets were then merged using bedtools merge into a single file and only elements between 20 bp and 5000 bp were retained. The UCSC dataset was originally in the hg38 human coordinates. We used UCSCs *liftover* tool using the chain file hg38TogalGal4 to first convert to chicken galGal4 coordinate system, and then subsequently to galGal7b coordinates system using NCBI Genome Remapping Service. Similar to the other two datasets, fragmented regions were merged together, and then elements less than 20 bp and greater than 5000 bp were filtered out.

To generate a new list of conserved elements de novo, we used phastCons ^78^. Before we can get a list of conserved elements, we need to calculate the rates of neutral evolution and conserved evolution. For neutral rates, we generated a bed file of four-fold degenerated sites from our 79-genome cactus alignment using the *hal4dExtract* command of the HALtools package and the annotated feature files (gff) for chicken. We generated separate bed files for chicken macrochromosomes (1 through 12), microchromosomes (13 through 38), and Z chromosome. Alignments for the four-fold degenerate sites were generated using the *hal2maf* command of the HALtools package, with galGal7b as the reference genome. The neutral rate for macrochromosomes, microchromosomes, and the Z chromosome was then estimated using phyloFit ^79^. For the conserved rate, we generated whole chromosome alignments for only non-target lineages since we are interested in acceleration in conserved sites in target lineages. We ran phastCons to estimate conserved rate by estimating ρ (rho), the mean substitution rate in conserved elements relative to the neutral rate. The values of ρ were estimated for each chicken chromosome independently. For macrochromosomes, we used a GC value of 0.42 (mean of GC content of all macrochromosomes), 0.52 for microchromosomes (mean of GC content of all microchromosomes), and 0.41 for the Z chromosome. The average conserved rate for macrochromosomes, microchromosomes, and the Z chromosome was then estimated using *phyloBoot*. Finally, we generated a list of conserved elements using phastCons with the --*most-conserved* flag. Elements smaller than 20 bp and larger than 5000 bp were subsequently filtered out.

We also generated a second set of conserved elements using the zebrafinch, *Taeniopygia guttata*, genome as the reference to get a list of elements that were not biased to the chicken genome. We used the same neutral rate and conserved rates generated from above to run phastCons with the alignments referenced on the *T. guttata* genome. We considered *T. guttata* chromosomes 1 through 12, including 1A and 4A, as macrochromosomes, and the remaining non-Z chromosomes and scaffolds as microchromosomes. We used *halLiftOver* command of the HALtools package to lift over the *T. guttata* coordinate system to the galGal7b coordinate system. Fragmented regions were merged with a gap tolerance (-d) of 5 bp, and then elements less than 20 bp and greater than 5000 bp were filtered out.

All five datasets were merged into one and elements less than 20 bp and greater than 5000 bp were filtered out from this combined dataset. The dataset was annotated for which of the five datasets the element included the element (Supp. Fig. 16), whether the conserved elements were located in exons, introns, or intergenic regions, whether they were located in known ATAC-seq peaks for chicken (see below for methods for ATAC seq), what fraction of the bases were considered conserved in the per-base conservation score from Feng et al. (2020) for 363-way alignment of bird genomes, and phyloP conservation scores. For phyloP conservation scores, we used alignments of each conserved element and then used the CONACC mode using an LRT method implement in phyloP. Elements with phyloP scale scores greater than 20 were omitted from further analysis. We also filtered elements that had both high phyloP conservation scores (>0.6) and low fraction of sites conserved (<0.3). This dataset constitutes the final dataset of conserved elements.

To compare the avian elements to those in mammals to assess patterns of convergence, we projected the mammalian phyloP conservation scores from the Zoonomia mammalian alignment ^19^. We converted the bigwig file to wig format using UCSC *bigWigToWig* program. Using a q-value of 0.05 after performing a Benjamini-Hochberg (BH) correction ^80^ on the PhyloP p-values, we extracted the list of sites that are conserved in mammals. These sites were then converted from human coordinates (hg38) to chicken coordinates (galGal7b) using the tool liftOver. Overlap of successfully lifted over sites with the avian conserved elements was assessed using bedtools intersect function. Elements that were not lifted over were labelled as NA while those elements with conserved fractions greater than 0.3 were assumed to be conserved in both mammals and birds.

#### Generating list of ATAC-seq peaks

Along with a data set of conserved elements, we also gathered a list of ATAC-seq peaks to delineate elements that are in open chromatin regions versus closed chromatin regions. For this we gathered published chicken ATAC-seq files from eleven studies encompassing 36 tissue types from different embryonic developmental stages that were uploaded to the SRA database (Supp. Table 4). Altogether, we had 101 fastq files including replicates for each tissue type. We used NGmerge (https://github.com/jsh58/NGmerge) to remove adapters and clean up fastq files and then used bwa v0.7.17 ^81^ to map each fastq file to the galGal7b reference genome. We used Genrich (https://github.com/jsh58/Genrich) to call ATAC-seq peaks. We ran Genrich in ATAC-seq mode with PCR duplicate removal mode for each tissue type and also one run using all tissue types together. Using all files to call peaks facilitates confident calling of peaks present in multiple tissue types. We then used bedtools intersect with the -v flag to get peaks that are unique to each tissue type and not present in the combined Genrich call. Finally, we merged all the unique elements to generate the final ATAC-seq bed file.

We annotate the ATAC-seq bed file using bedtools *annotate* to create a matrix of presence/absence of ATAC-seq peak in any particular tissue. To visualize the relationships of the ATAC-seq peaks we used the presence/absence matrix as an input for iqtree2 ^82^ with a MK model with ascertainment bias to generate a tree.

#### Running PhyloAcc

We filtered our list of conserved elements to those not occurring only in exons using bedtools intersect resulting in a list of conserved non-exonic elements (CNEE). For CNEES in each chromosome, we generated a maf alignment using hal2maf from HALtools with *Gallus gallus7* as the reference genome. We used the *--noDupes* flag to output only a single sequence for each species in case there were paralogs and *--noAncestors* flags to only include sequences from the tips. For each maf alignment, we used a custom script to generate fasta alignments for each CNEE (https://github.com/subirshakya/phast_scripts). We then used another custom script to concatenate the fasta alignments of all macrochromosomes, all microchromosomes, and all Z chromosomes CNEEs. Here we also filtered any elements that were missing more than 10% of taxa (Supp. Fig. 17). We omitted the elements from the W chromosome due to the absence of W chromosomes in many of the genomes used. We used the concatenated alignment and the neutral rates generated above as input for PhyloAcc ^33^. Three instances of PhyloAcc were run using the species tree mode for macrochromosomes, microchromosomes, and Z chromosomes. All species for clades with shifts to short tarsus length were used as target species. *Gavialis gangeticus* was set as outgroup taxon.

We subset the full dataset based on Bayes factor of 10 or greater for the comparison between the null model (with no acceleration allowed), and the target-lineage model (where acceleration is allowed in only the focal/target lineages). PhyloAcc includes an assumption of Dollo’s irreversible evolution hypothesis ^33,83,84^ as such a shift to acceleration at one branch means all subsequent child branches for that node will also be in accelerated state. Hence, we consider acceleration at the base of target clade as a sign of acceleration of all focal taxa within that node. We used this to retain elements where all members of the accelerated focal clade (or more than 3 penguins) are accelerated (short-tarsi broad dataset). Subsequently we identified a subset of accelerated elements where the Bayes factor for the comparison between the target-lineage model (where acceleration is allowed in only the focal/target lineages) and the unrestricted model (where acceleration can occur anywhere) was at least 5, indicating more support for the target-lineage than the unrestricted model (short-tarsi specific dataset).

#### Changes in ATAC-seq peaks

While we can identify elements that overlap with ATAC-seq peaks from chicken, we do not have information whether a region is in open or closed chromatin regions in other species. One approach to predict open chromatin region in disparate species using known ATAC-seq profiles is TACIT, a machine learning framework developed by Kaplow et al. (2022). We used open regions from Hindlimb HH18 stage in chicken and emu, *Dromaius novaehollandiae,* ^44^ for our positive dataset. For emu, we mapped the fastq files to the emu genome, droNov1, using the methods described above for chicken. For all datasets we used sequences that were 500 bp long measured from the center of the peak for each ATAC-seq peak. We only selected one peak if two peaks were within 500 bp of one another to prevent counting the same peak twice. For the negative dataset, we used peaks that are present in chicken but absent in emu and vice versa. For this, we used *halLiftOver* command of the HALtools package to lift over the chicken coordinate system to emu coordinate system and vice versa for the respective fasta files of ATAC-seq peaks. Fragments that are within 100 bp were merged in the lifted over files. We then kept elements that were between 400-600 bp and then reformatted them to be 500 bp long by selecting the center of the sequence and extending out. We then used bedtools intersect to get peaks in chicken that do not overlap with emu peaks and similarly, emu peaks that do not overlap with chicken peaks. We combined these two to get the negative dataset. We filtered out any peak in both the positive and negative dataset that overlapped with coding regions.

For training, validation, and evaluation in the machine learning framework, we divided the dataset into three parts based on the size of the chromosome. The training dataset contained elements from the largest two chromosomes in both chicken and emu. The validation dataset contained elements from the third largest chromosome and then the evaluation dataset contained elements from the fourth largest chromosomes. We then cycled through the remaining chromosomes in the same pattern. This was done for both the positive and the negative dataset.

We then used the code from ^20^ to train the model. We used five convolution layers with 450 filters of width 7, stride 1, L2 regularization of 1e^-7^, and dropout of 0.2 followed by a max-pooling layer with 300 filters with width and stride 26. The model used was stochastic gradient descent with a cyclic learning rate and momentum, where the learning rate range was 0.00001 to 0.01, the momentum range was 0.90 to 0.99, and the number of cycles was 2.5. We used a batch size of 128 and we ran the model for 20 epochs, while checking for convergence.

Once we had our trained model, we then generated fasta files for all accelerated CNEEs for all species in our alignment. For this, we created a fasta file of the center position of each accelerated element for chicken and then used *halLiftOver* to convert this position to the coordinate system of every species in our alignment. Then we extended the position of each position to create a 500 bp region and extracted the 500 bp region from the respective genomes. We combined all these sequences to use as input for evaluation using our model. We get a probability of being present in an open-chromatin region for each CNEE for each species. We then compared the probability of being in a peak in an accelerated lineage to those in non-accelerated lineages to find elements where there have been changes in open chromatin profile that correlated with acceleration.

#### GO statistical overrepresentation test

We identified the genes 10,000 bp upstream and downstream of each element to get a list of genes around accelerated elements. This list contains genes that are potentially regulated by the elements around them. We used PANTHER ^85^ to perform a statistical overrepresentation test of genes around accelerated elements. We used the dataset biological process dataset (GO:0008150) for *Gallus gallus* (organism 9031) for GO classification. Since the genes around CNEEs are not randomly represented throughout the genome, to create a baseline approximation for overrepresentation, we generated a list of all genes near all CNEEs for the reference list. We did 1000 iterations where we subsampled the same number of CNEEs from the full CNEE dataset as the test state and used the genes around the elements. We then compared the observed value for number of genes in each pathway to the distribution of 10,000 random repeats to calculate and approximate p-value of significance and calculate enrichment scores. GO categories with a p-value less than 0.05 were considered overrepresented. This process was done for both the full dataset and ingroup dataset. Since most GO categories contain overlapping sets of genes, we used a Gene Ontology (GO) semantic similarity matrix to categorize overrepresented categories into clusters. This was done by creating a similarity matrix using *GO_similarity* function of the simplifyEnrichment package ^45^ in R. We then used the *simplifyGO* to cluster the GO categories using a binary cut clustering algorithm ^45^.

#### Test for convergence in accelerated elements

To test for convergence in accelerated elements, we performed pairwise hypergeometric tests for each group of short-tarsi focal clades. We also selected four representative outgroups, which have proportionally similar total branch lengths within the clade and are relatively equidistant to the other ingroup clades as the representative ingroup clade. For bulbuls, we used the clade containing *Sturnus vulgaris* and *Ficedula albicollis*; for swallows, we used clade containing *Emberiza elegans*, *Molothrus ater*, *Setophaga coronata*, and *Diglossa brunneiventris*; for kingfishers, we used clade containing *Pogoniulus pusillus* and *Colaptes auratus*. We do not have any clade with similar taxonomic breath as penguins, so we used a clade containing only two species *Sterna hirundo* and *Pluvialis apricaria*, while acknowledging the shortfalls. To get a list of accelerated elements in the outgroup clades, we first subset the full dataset based on Bayes factor of 10 or greater for the comparison between the null model (with no acceleration allowed), and the full model (where acceleration is allowed in only all lineages). We then filtered this set of accelerated elements to retain elements where all members of the accelerated focal outgroup clade are accelerated. We then used Fisher’s exact test to compare the number of accelerated and convergent elements in the focal groups with those of the outgroups to estimate whether there is an increased number of accelerated elements in the focal groups. We performed six pairwise tests, four three-way comparisons, and one four-way comparison.

#### Hindlimb development and changes in motifs sequences

To create a list of genes associated with hindlimb development, we used the schemes in ^43^ for early embryonic development of limb buds and bone development. For each gene, we identified elements near the representative genes and categorized them based on presence in one of the four target lineages. Next, we used the JASPAR vertebrates CORE non-redundant motif file ^86^ to find motifs that correspond to the genes identified for hindlimb development. We then used me MAST program in the MEME Suite v5.2.2 package, using default settings, to locate the motifs in the alignments for the identified elements ^87,88^. We then looked for elements where acceleration and motif gain/loss are correlated.

#### Positive selection analysis

We also tested for positive selection in lineages associated with short taxa to contrast with results from acceleration in regulatory CNEEs. To get an alignment of protein coding loci we used TOGA ^15^ to annotate all 77 genomes, excluding the two chicken genomes, using the annotation of the galGal7b chicken genome. TOGA uses a machine learning and synteny based approach to infer orthologous regions of the genome. For every species, we used a chain file between that species and the chicken, that was extracted from the 79-way cactus alignment.

Briefly, we used halLiftOver to convert the alignment to a psl file. We then forced positive strandedness in the psl file using pslPosTarget (UCSC). Finally use used axtChain (UCSC) to convert the psl file to a chain file. This chain file and all the annotated coding sequences of the chicken genome were used as inputs for TOGA.

We used the TOGA outputs to generate a list of genes labelled one2one, one-to-one orthologs, from all 77 genomes. We also included one-to-one genes where in some taxa the gene was not found, labelled one2zero. Similarly for species with genes labelled one-to-many, we excluded these from the final list of genes but kept all the other species with one-to-one orthology. We kept the largest transcript for each gene and dropped any gene with fewer than 15 taxa represented. We then used a codon-aware aligner, macse v2 ^89^ to align all the genes. We filtered bad codons and misaligned species using a filtering algorithm (https://github.com/gwct/murine-discordance/blob/main/scripts/03-selection-tests/08_aln_filter.py). We further processed the alignments by masking misaligned codons using HMMCleaner ^90^. We used the four cost parameters -0.25, -0.20, 0.15, 0.45 for HMMCleaner.

For assessing positive selection, we used aBS-REL v2.3 ^64^. We ran aBS-REL while accounting for variable synonymous substitution rates ^91^, with a guide tree based with the topology based on the neutral tree used for PhyloAcc. We then filtered for genes with signs of positive selection in branches that included our focal clades with a corrected p-value that was significant. aBS-REL uses a Holm-Bonferroni correction ^92^ to correct for multiple tests. We also filtered out genes where there were fewer selected branches in the focal short -tarsi groups than outgroups. We then performed a GO statistical overrepresentation test as described previously to find genes in enriched GO categories. We repeated the same procedure for the outgroups (used previously for elements) and used Fischer’s exact test to test for convergence in selection in coding sequences.

#### McDonald and Kreitman tests for selection

We also used an imputed McDonald and Kreitman test for selection ^66^ to infer selection among short-tarsi lineages. For this, we only used bulbul and swallow as whole genome resequencing data is only available for species within those two lineages. For bulbuls, we used the population level resequencing data for *Brachypodius melanocephalos* from ^34^ and for swallows, we used data for *Hirundo rustica* from ^35^. We realigned all the individuals to the reference genome using snpArcher ^93^. For bulbuls, we subsequently dropped birds from small islands as the effects of genetic drift can affect subsequent results. Similarly, for swallows, we only used the individuals of the subspecies *rustica* to avoid issues from population structure and dropped one individual, SAMN18196751, due to the high degree of missing data. The following filters were applied to the variant call files (VCF) – kept only biallelic sites, indels were removed, removed sites with more than 25% individuals missing.

For the imputed MK test, we need polarized VCFs. To achieve this, we used an outgroup sequence to polarize all the SNPs within each vcf file. For bulbuls, we used *Sylvia atricapilla* (Genbank Ref: GCA_009819655.1) as outgroup, while for swallows, we used *Acrocephalus scripaceus* (Genbank Ref: GCA_910950805.1) as outgroup. We used the alignments from Cactus to generate a maf file for the whole genome alignment between the two species. We then converted this maf file into a vcf file using base by base comparisons between the two species. We filtered out indels and missing sites from the vcf file of the outgroup and then used bcftools merge to combine the outgroup vcf to the population resequencing vcf file. We then used vcfdo (https://github.com/IDEELResearch/vcfdo) to polarize the vcf files. We added a second outgroup to each vcf file for the MK test, *Phylloscopus trochilus* (Genbank Ref: GCA_016584745.1) for bulbul and *Eremophila alpestris* (Genbank Ref: GCA_009792885.1) for swallow. We then used degenotate (https://github.com/harvardinformatics/degenotate) to perform the imputed MK test.

For degenotate we used protein coding annotations identified by TOGA as input. We only kept the largest transcript of each gene for the MK test. We then performed a GO statistical overrepresentation test as described previously to find positively selected genes in enriched GO categories for each species. We also did a GO statistical overrepresentation test for all positively selected genes from both bulbul and swallow combined.

#### MK test on elements

We also implemented a modified version of the MK test to test for selection in the conserved elements. For this, we calculated the number of fixed and polymorphic sites for each element and then compared these to the number of fixed and polymorphic sites of the synonymous bases in the coding sequence of the nearest three genes. We calculated direction of selection and p-value for each element and compared those values for accelerated elements in bulbuls and swallows respectively. For positively selected elements, we performed a GO enrichment analysis for genes that are within 10,000 bp of these elements, using the approach described in the previous section for elements.

We also performed a selection test for each gene based on the conserved element environment around the gene. For this, we selected all elements within 10,000 bp of each gene, including elements contained within the intronic regions of the gene. We filtered out elements that were accelerated in the outgroup branch to omit a potential source of false positives. We then summed up the number of fixed and polymorphic sites of each of the conserved elements and calculated a p-value, direction of selection, and alpha for each gene. We selected genes that were positively selected with a p-value less than 0.0005, with a positive direction of selection, and an alpha greater than 0.5. We performed GO enrichment analysis for this set of genes for bulbuls and swallows respectively.

## Data availability

Genome alignments, ATAC-seq calls, TOGA annotations, selection results, and all other relevant data will be made available in X after acceptance of manuscript.

## Code availability

All code for this study is available as a fully documented Rmarkdown file in https://github.com/subirshakya/projects_rmarkdown

## Acknowledgements

We thank Cliff Tabin, Patrick Gemmell, Emma Farley, Meng Zhu, and Granton Jindal for their comments on improving the manuscript. This project was supported by the NIH grant NHGRI R01HG011485. Computational resources for this project were provided by the FASRC Cannon cluster at Harvard University. We also thank Jeremiah Trimble and Kate Eldridge for access to the collections at the Museum of Comparative Zoology at Harvard University.

