## Supplementary Figures for "Convergent evolution of noncoding elements associated with short tarsus length in birds"

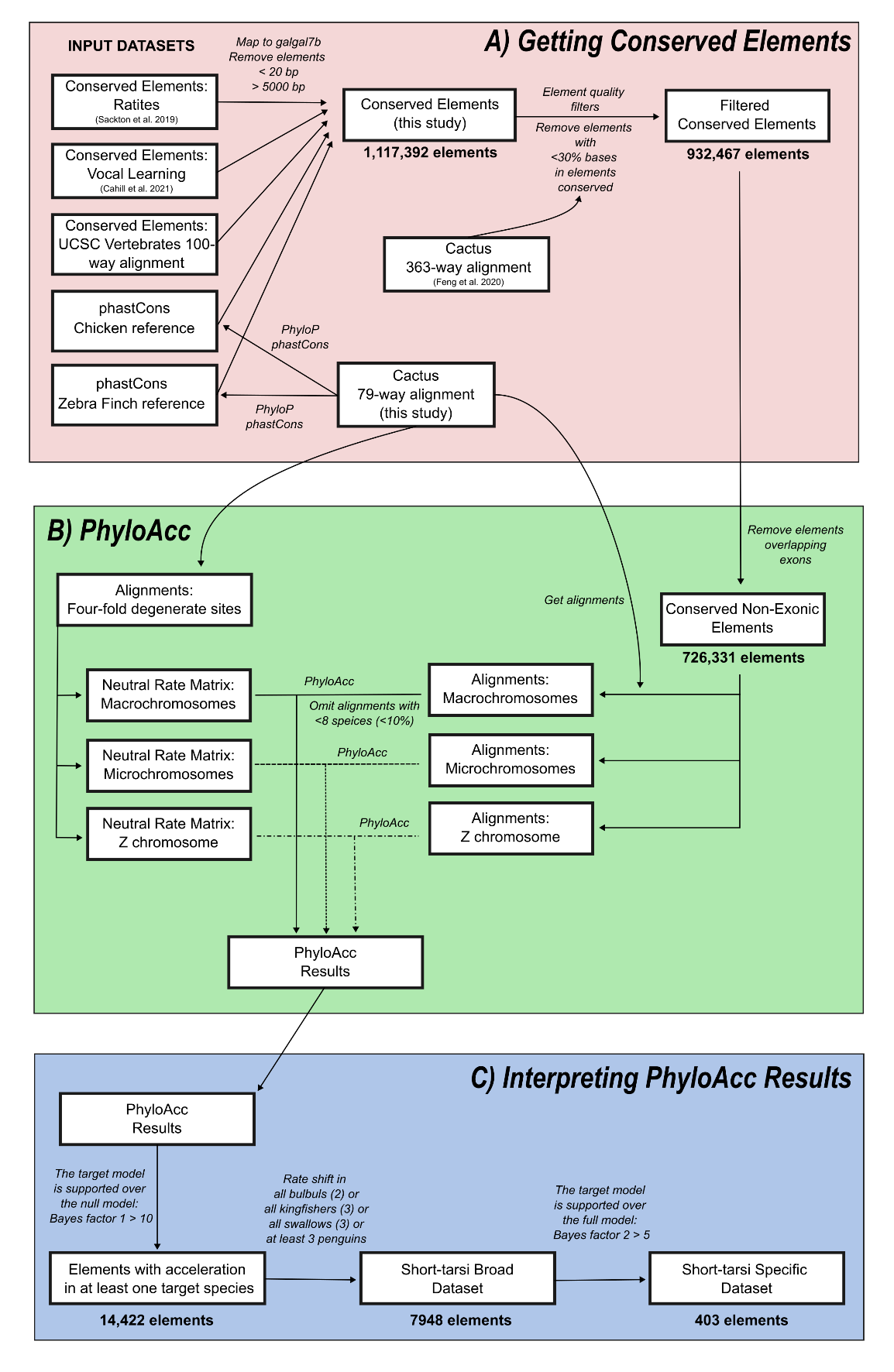

Supplementary Figure 1: Schematic of methods and datasets used to A) get conserved elements, B) run PhyloAcc, and C) interpret PhyloAcc results. Number of elements at different points in the schematic are represented below representative blocks.

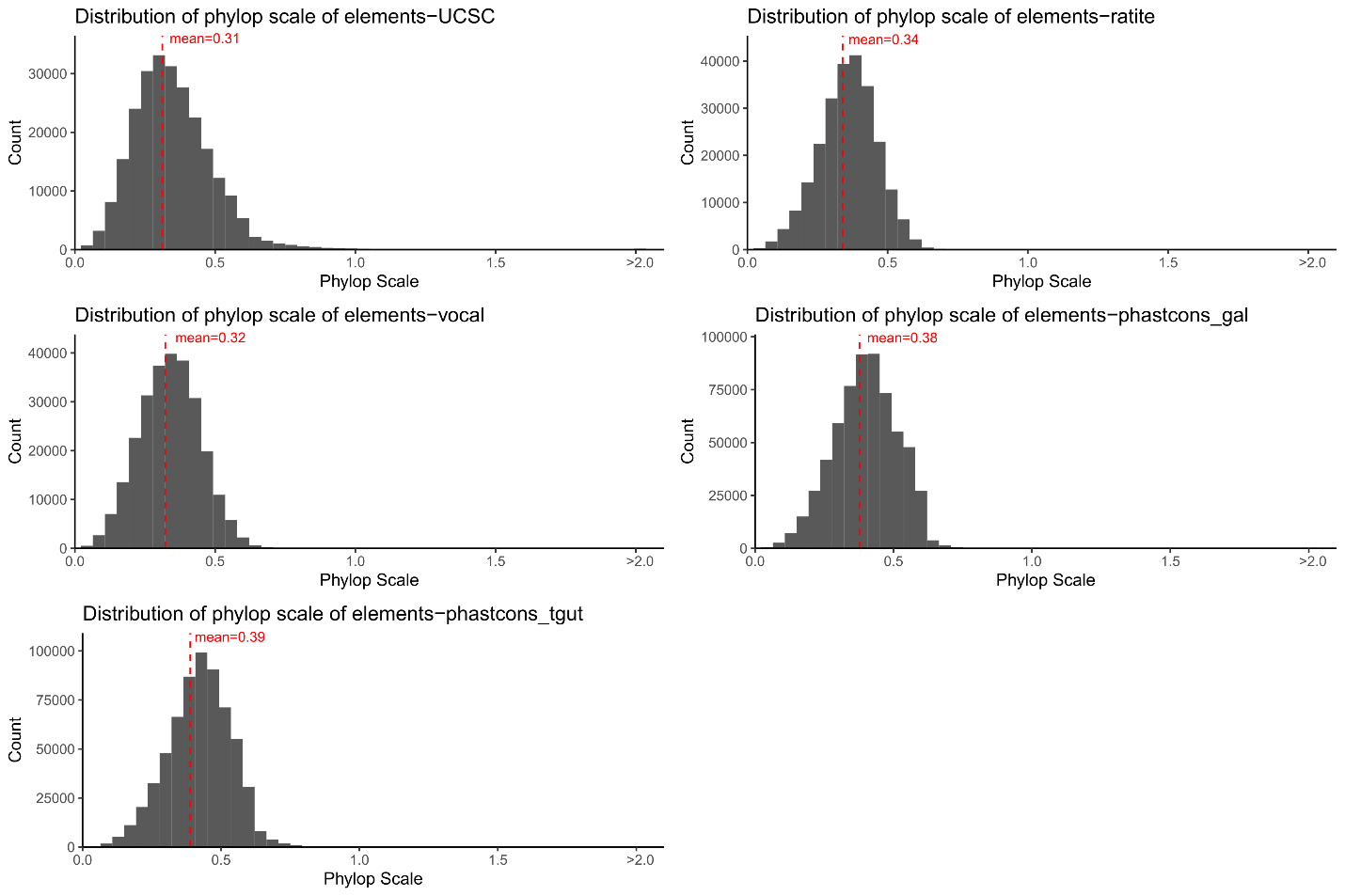

Supplementary Figure 2: Distribution of phylop scale values in the five source datasets: UCSC vertebrate-wide elements (UCSC), conserved elements used by Sackton et al., (2019) (ratite), conserved element used by Cahill et al., (2021) (vocal), conserved elements called using phastcons from the 79-way cactus alignment in this project with chicken as reference species (phastcons_gal) and zebra finch as reference species (phastcons_tgut). The weighted mean of each distribution is given by red line. Values greater than 2.0 have been collated into a single bin.

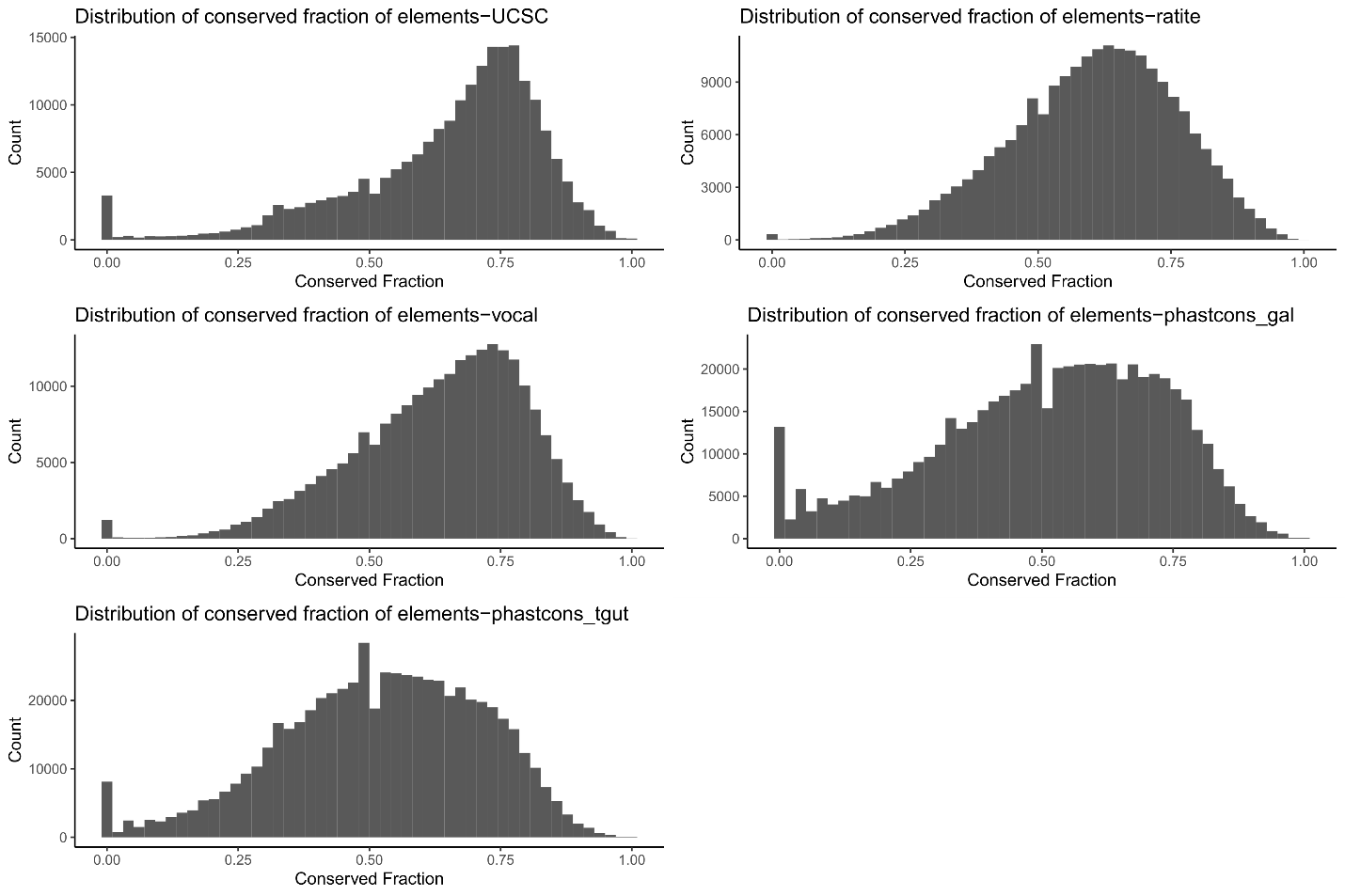

Supplementary Figure 3: Distribution of fraction of conservation in the five source datasets: UCSC vertebrate-wide elements (UCSC), conserved elements used by Sackton et al., (2019) (ratite), conserved element used by Cahill et al., (2021) (vocal), conserved elements called using phastcons from the 79-way cactus alignment in this project with chicken as reference species (phastcons_gal) and zebra finch as reference species (phastcons_tgut).

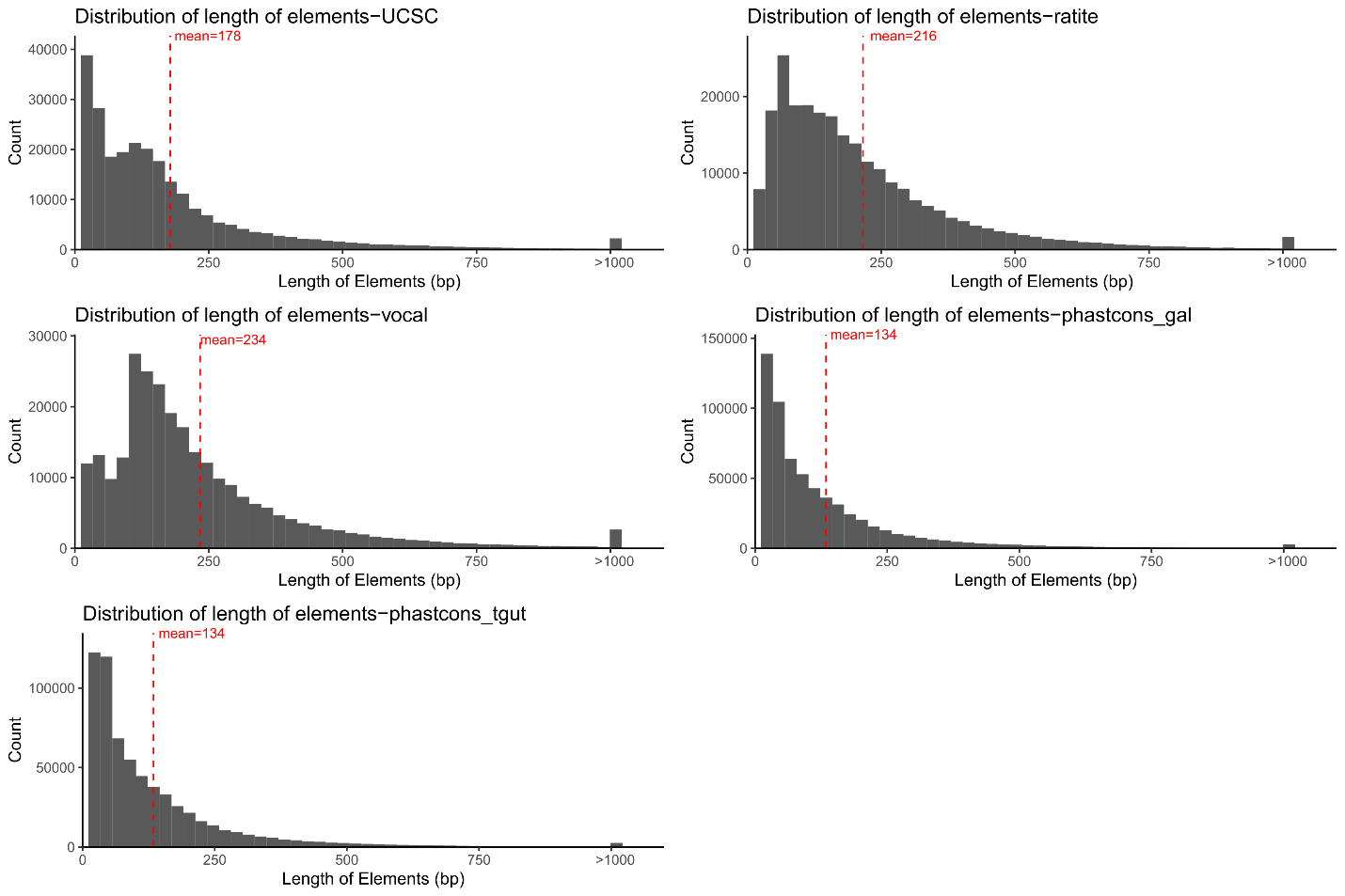

Supplementary Figure 4: Distribution of length of conserved element divided by the five source datasets: UCSC vertebrate-wide elements (UCSC), conserved elements used by Sackton et al., (2019) (ratite), conserved element used by Cahill et al., (2021) (vocal), conserved elements called using phastcons from the 79-way cactus alignment in this project with chicken as reference species (phastcons_gal) and zebra finch as reference species (phastcons_tgut). The mean lengths of the elements are shown by red line. Elements larger than 1000 bp have been collated into a single bin.

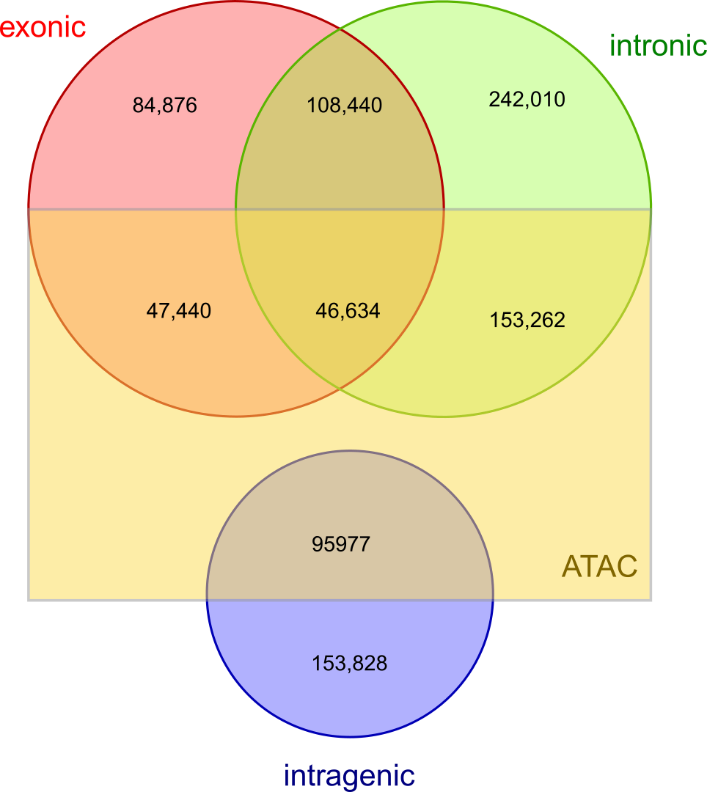

Supplementary Figure 5: Venn diagram showing overlap of conserved elements with exons, introns, and intergenic regions. Regions that overlap ATAC-seq peaks are delineated in the box.

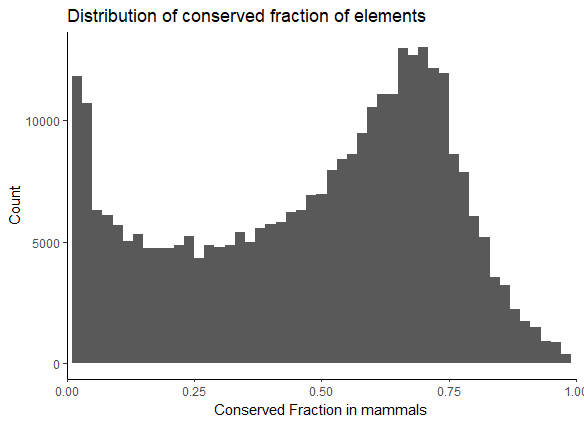

Supplementary Figure 6: Distribution of fraction of conservation among conserved elements in birds with respect to overlapping regions in the Zoonomia mammal alignment (Armstrong et al., 2020).

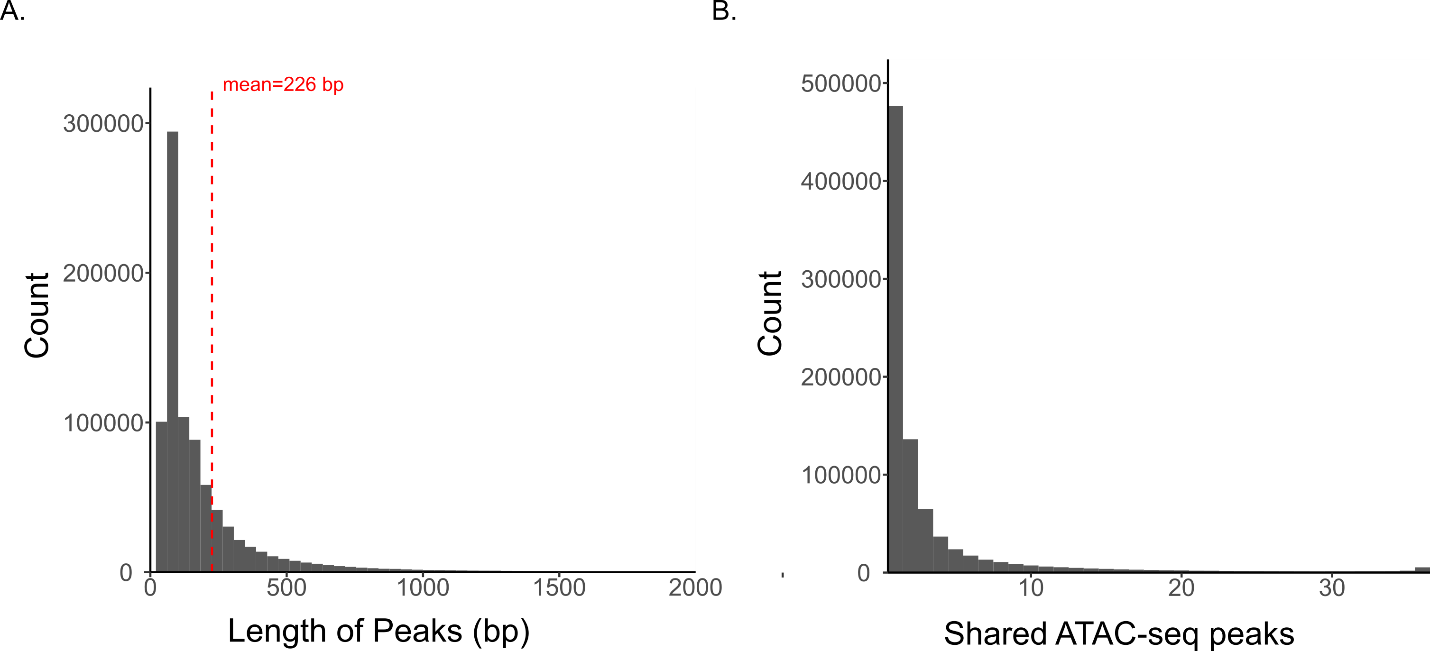

Supplementary Figure 7: A) Distribution of lengths of ATAC-seq peaks called from 36 embryonic tissues at different stages for chicken. Peaks greater than 2000 bp have been collated into one stack. The red line shows the mean length of the peaks. B) Distribution of shared tissues for each ATAC-seq peak going from one, present in only one tissue, to 36, present in all 36 tissues for which we called ATAC-seq peaks.

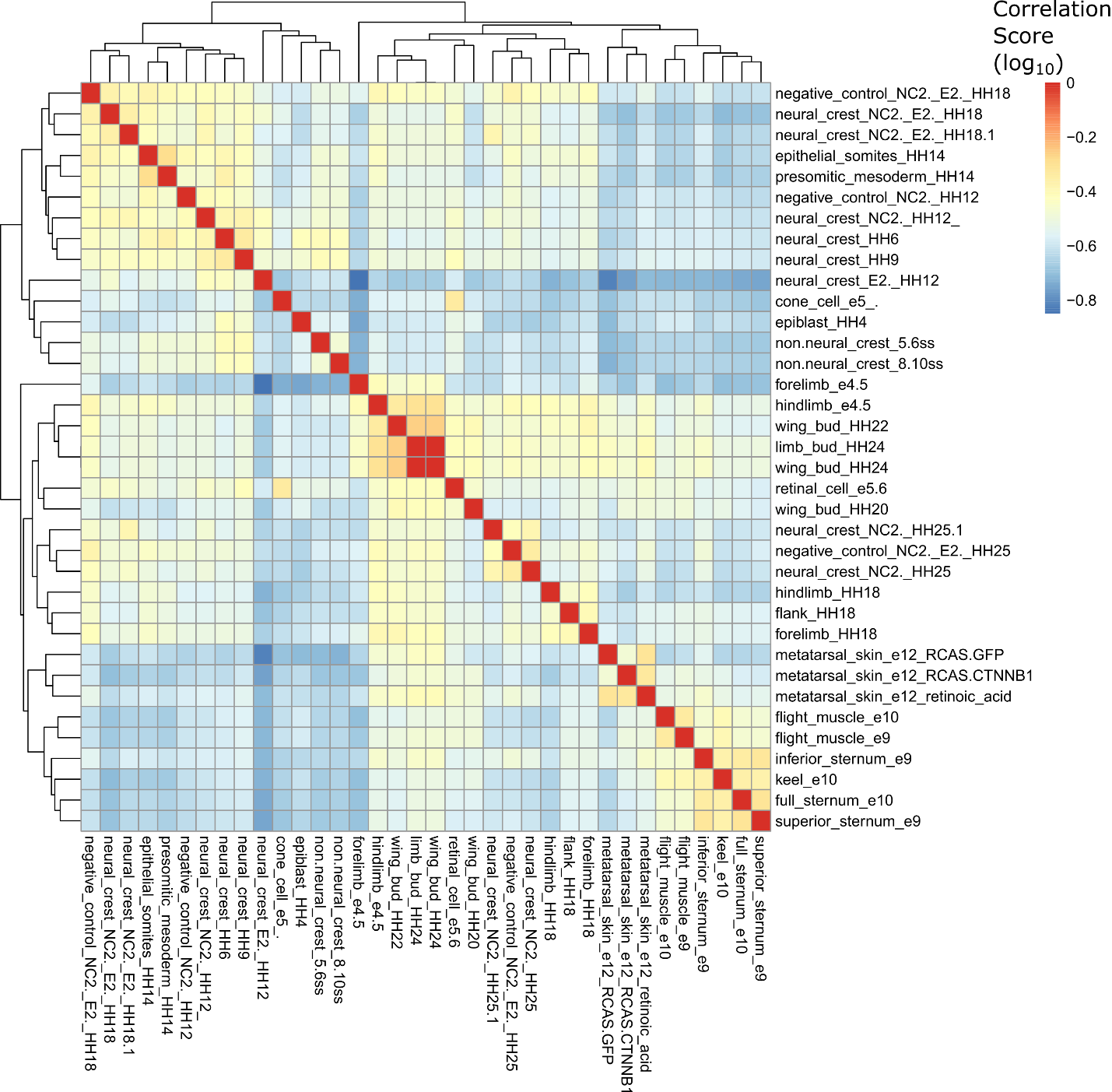

Supplementary Figure 8: Correlation heat-map showing relationship among different tissue types based on shared ATAC-seq peaks. Correlation values have been log transformed.

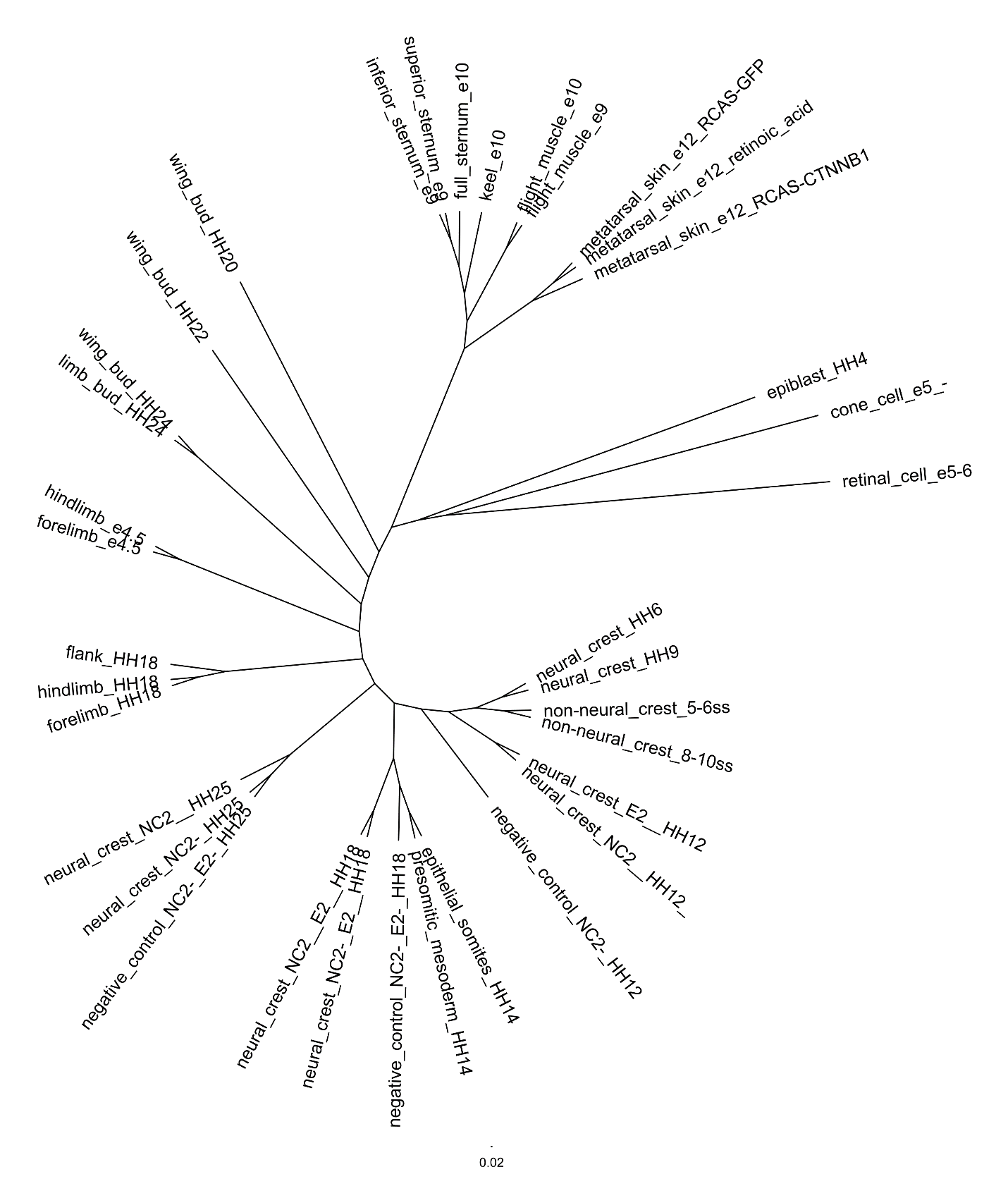

Supplementary Figure 9: Tree diagram showing relationships between different tissues from ATAC-seq peaks at different embryonic stages of chicken. The tree is based on a presence/absence matrix of peaks and is inferred using a Maximum Likelihood framework in IQtree2.

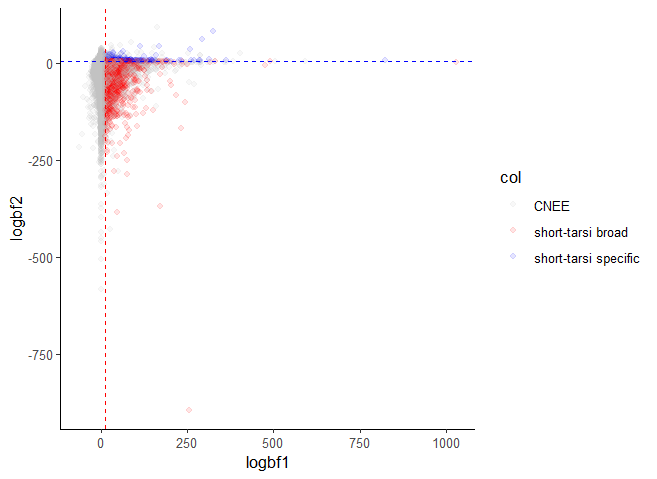

Supplementary Figure 10: Plot of log of Bayes Factor 1, for the comparison between the null model (with no acceleration allowed) and Bayes Factor 2, for the comparison between the target-lineage model (where acceleration is only allowed in only the focal/target lineages) and the unrestricted model (where acceleration can occur anywhere). The red line denotes Bayes Factor 1 of 10 and points to the right of this line constitutes the short-tarsi broad dataset of accelerated elements. The blue line denotes Bayes Factor 2 of 5 and points above this line and to the right of the Bayes Factor 1 cutoff constitutes the short-tarsi specific dataset of accelerated elements.

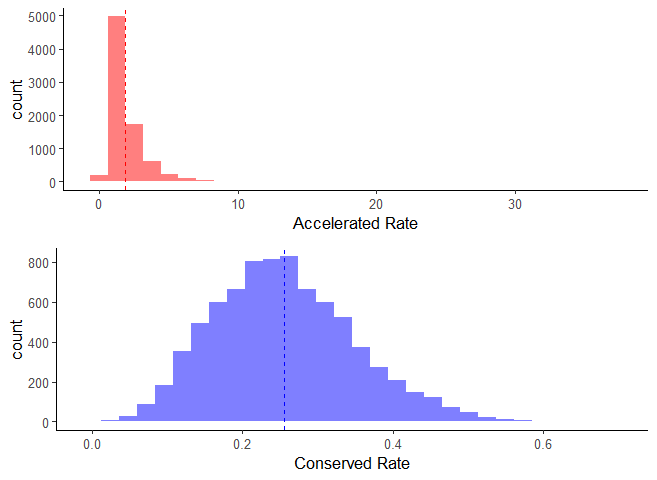

Supplementary Figure 11: Histogram showing distribution of accelerated and conserved rates for elements in the short-tarsi broad dataset. The dotted line shows the mean rate for each distribution.

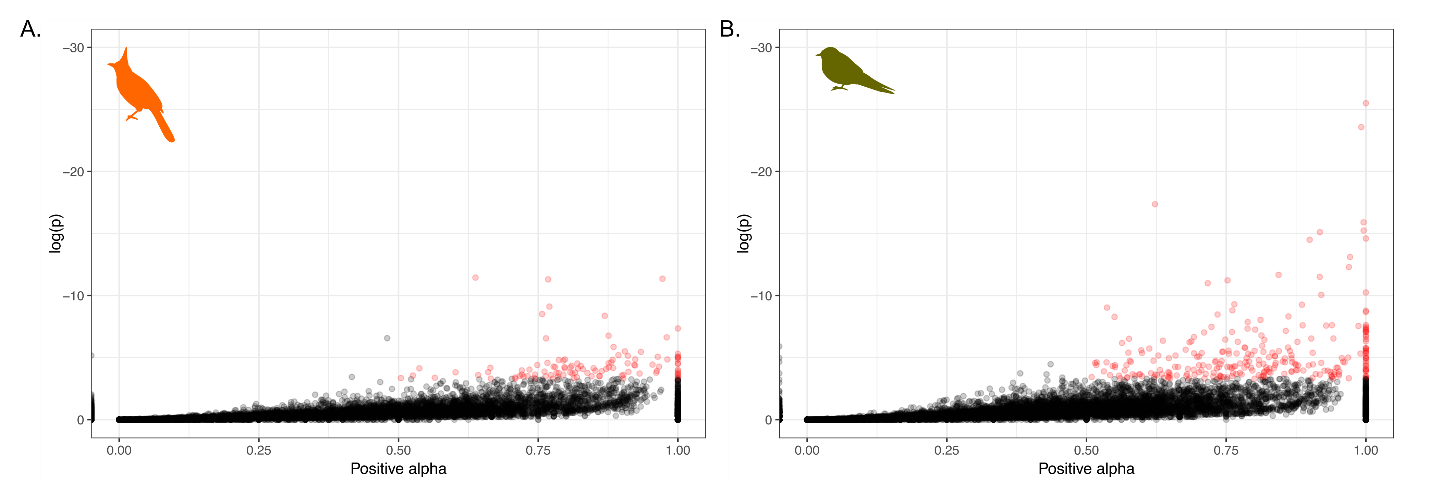

Supplementary Figure 12: Significant alpha (positive selection) values from a modified McDonald & Kreitman test using fixed and polymorphic sites of all conserved elements within 10kb of a gene compared to fixed and polymorphic sites in synonymous sites in the gene for A) bulbuls and B) swallows.

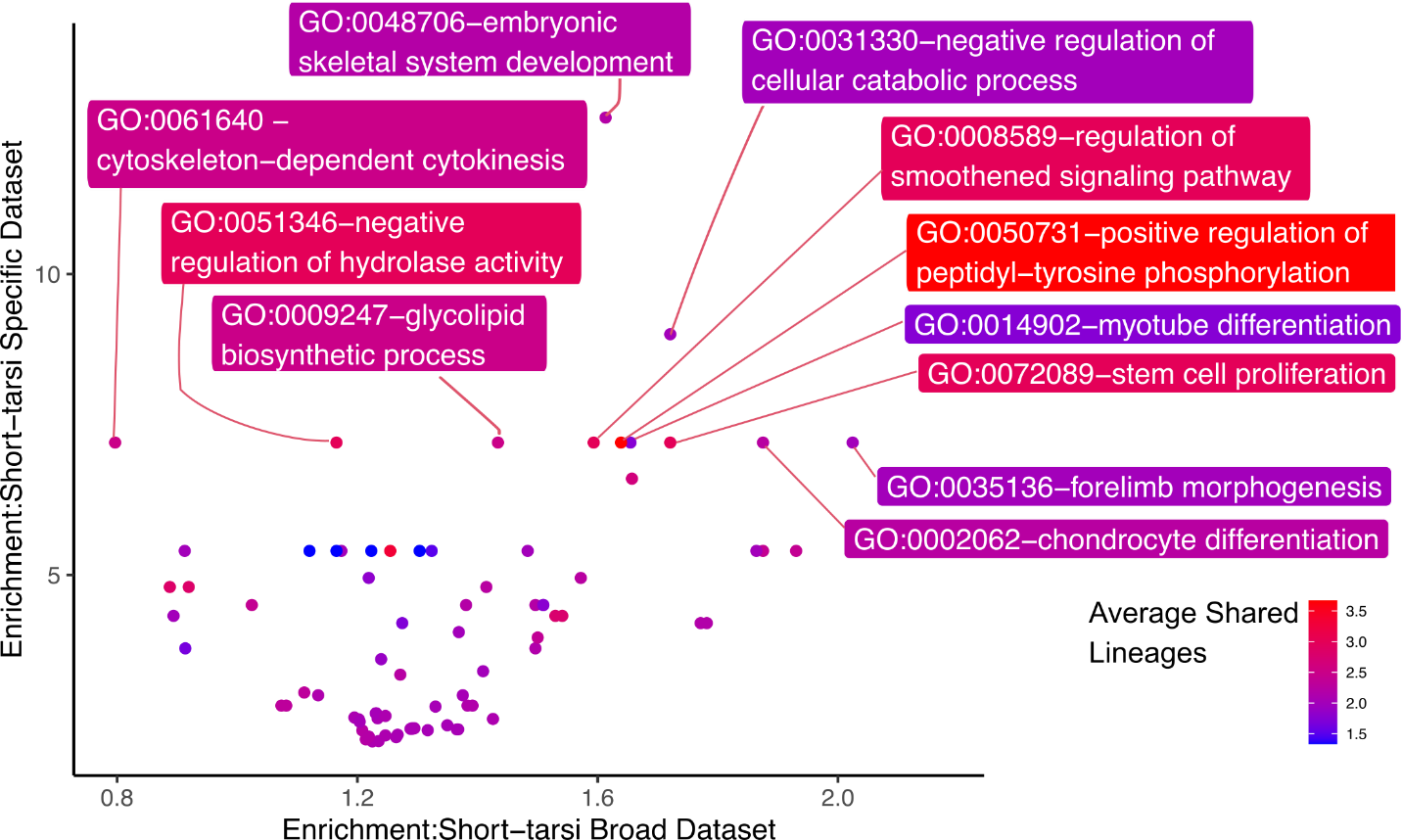

Supplementary Figure 13: Scatterplot between enrichment scores for GO categories in the full dataset and ingroup dataset. Points are colored based on the average number of focal clades showing acceleration in CNEEs constituting genes in that GO category.

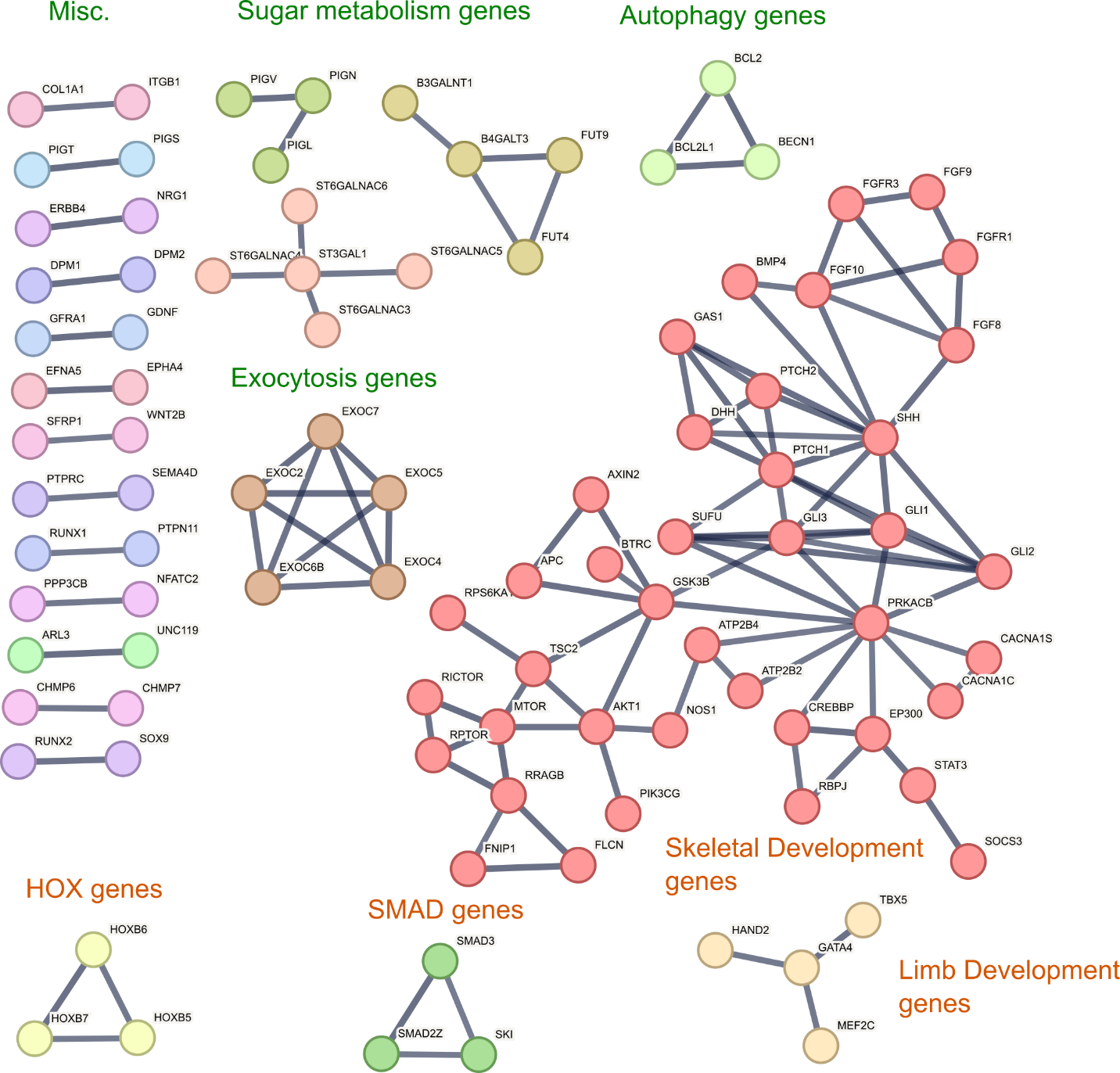

Supplementary Figure 14: STRING network graph of genes that are near accelerated elements are listed significantly enriched GO categories in the short-tarsi specific dataset when compared to the short-tarsi broad dataset. Each circle represents a gene with the lines connecting them representing high confidence (0.9) known and predicted interactions between the genes from STRING database. Major networks are colored by groups.

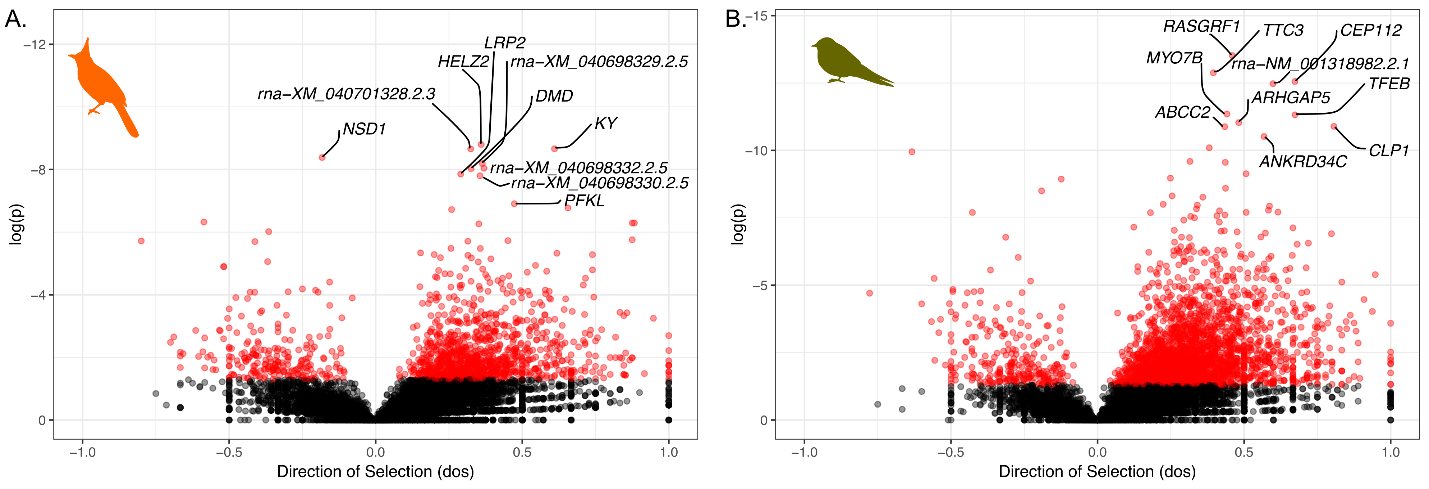

Supplementary Figure 15: Direction of selection from an imputed McDonald & Kreitman test for genes in A) bulbuls and B) swallows. Significant values are represented in red.

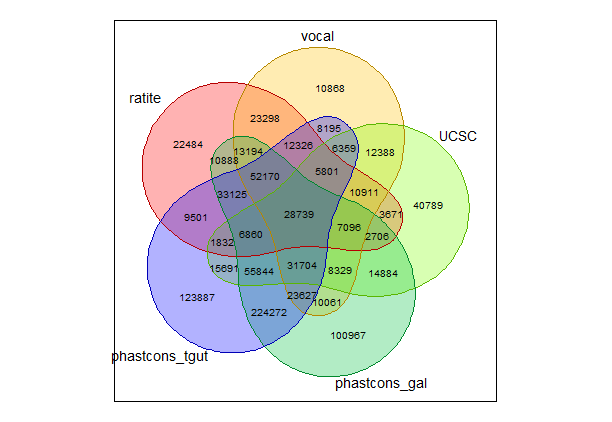

Supplementary Figure 16: Venn-diagram showing the elements shared between the five source datasets: UCSC vertebrate-wide elements (UCSC), conserved elements used by Sackton et al., (2019) (ratite), conserved element used by Cahill et al., (2021) (vocal), conserved elements called using phastcons from the 79-way cactus alignment in this project with chicken as reference species (phastcons_gal) and zebra finch as reference species (phastcons_tgut).

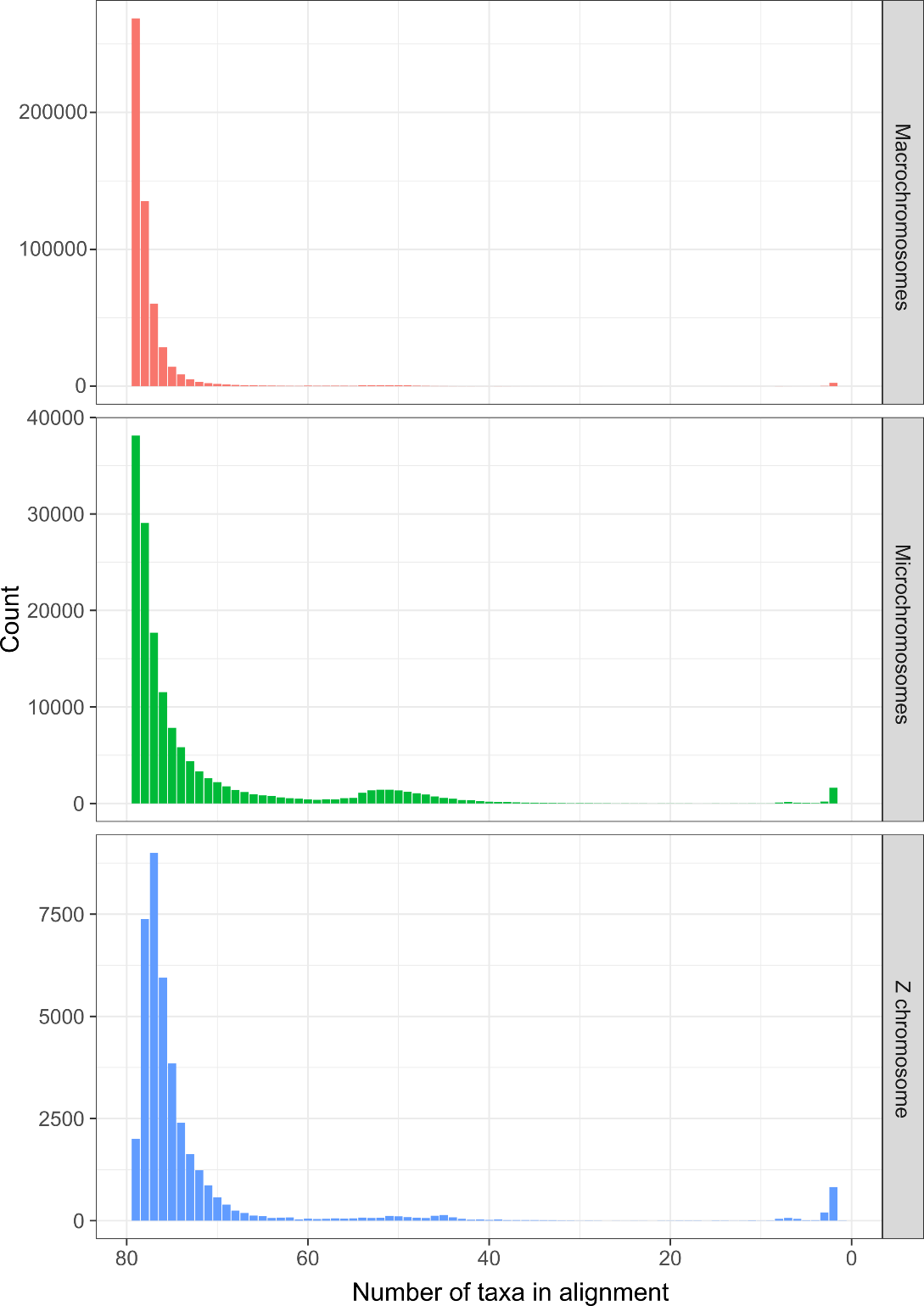

Supplementary Figure 17: Distribution of the number of taxa present in alignments of each CNEE separated by macrochromosomes, microchromosomes, and Z chromosome.

**Supplementary Tables:**

Supplementary Table 1: A) Run statistics for two independent runs for each of the fourteen groups in bayou. The table notes the estimated values, effective sample sizes, and Gelman’s R statistic for each of log likelihood (lnL), prior density, alpha parameter, sigma-squared parameter, number of shifts (n), and number of optima (theta). B) Taxa showing shifts in tarsus length size detected from bayou and the posterior probabilities and theta supporting shifts of each group from the two runs. C) Subset of AVONET dataset showing data used for this study and the assignment to the fourteen groups used in this study.

Supplementary Table 2: Measurements of femur, tibiotarsus, and tarsometatarsus for specimens of the four focal groups (penguins, kingfishers, bulbuls, and swallows) and their outgroups. The respective tarsus length and mass from AVONET for each species is also listed.

Supplementary Table 3: Accession Numbers and basic genomic statistics for genomes used in the 79-way Cactus alignment. Genome Statistics are reported from NCBI for genomes downloaded from NCBI.

Supplementary Table 4: List of datasets used in the generation of ATACseq panel. SRA accession numbers for each replicate from corresponding studies are listed in the SRA accession column. The total reads mapped shows the number of reads mapped to the galGal7b genome. Citations for respective studies used are at the bottom of the table.

Supplementary Table 5: Table showing predicted ATAC-seq peaks for accelerated elements using the trained CNN model for all accelerated elements compared to the true overlap of peaks from sequencing data for Hindlimb HH18 stage in chicken*.* Values in parenthesis with * show elements which share the same predicted state in 50% of the accelerated taxa for that element from PhyloAcc.

|  | | Real ATAC-seq peaks for chicken | |
| --- | --- | --- | --- |
|  |  | TRUE | FALSE |
| Predicted ATAC-seq peaks for chicken using trained model in TACIT | TRUE | 256 (146*) | 1155 (658*) |
|  | FALSE | 1113 (1096*) | 5064 (4965*) |

Supplementary Table 6: GO (Gene Ontology) terms and fold enrichment for genes near accelerated elements in the A) short-tarsi broad and B) short-tarsi specific datasets.

Supplementary Table 7: Test of convergence for shared positively selected genes from branch-site tests of selection between four focal taxa. Pairwise hypergeometric tests were done for the four focal groups, and Fisher’s exact test were done using representative outgroups for each group. Significant values are represented in bold. The Fisher’s exact test for all four groups could not be performed as there were no genes with positive selection shared by all four groups.

| Bulbul | Swallow | Kingfisher | Penguin | Hypergeometric test p-value | Fisher’s exact test p-value | Odds ratio |
| --- | --- | --- | --- | --- | --- | --- |
| X | X | X | X |  | NA | NA |
| X | X | X |  |  | 1 | 0 |
| X | X |  | X |  | 0.74 | 0.98 |
| X |  | X | X |  | 0.95 | 0.98 |
|  | X | X | X |  | 1 | 0 |
| X | X |  |  | 0.52 | 1 | 0.07 |
| X |  | X |  | 1 | 0.12 | 0.39 |
| X |  |  | X | **5.65** **x 10^-19^** | 0.68 | 0.92 |
|  | X | X |  | **2.32 x 10^-3^** | 0.99 | 0.09 |
|  | X |  | X | **1.82 x 10^-53^** | 0.29 | 0.95 |
|  |  | X | X | **8.34 x 10^-56^** | 0.09 | 2.38 |
